# A Genetically encoded BRET-based SARS-CoV-2 Mpro protease activity sensor

**DOI:** 10.1101/2022.01.31.478460

**Authors:** Anupriya M Geethakumari, Wesam S Ahmed, Saad Rasool, Asma Fatima, S.M. Nasir Uddin, Mustapha Aouida, Kabir H Biswas

## Abstract

The SARS-CoV-2 main protease, M^pro^, is critical for its replication and is an appealing target for designing anti-SARS-CoV-2 agents. In this regard, a number of assays have been developed based on its cleavage sequence preferences to monitor its activity. These include the usage of Fluorescence Resonance Energy Transfer (FRET)-based substrates in vitro and a FlipGFP reporter, one which fluoresces after M^pro^-mediated cleavage, in live cells. Here, we have engineered a pair of genetically encoded, Bioluminescence Resonance Energy Transfer (BRET)-based sensors for detecting SARS-CoV-2 M^pro^ proteolytic activity in living host cells as well as in vitro assays. The sensors were generated by sandwiching M^pro^ N-terminal autocleavage sites, either AVLQSGFR (short) or KTSAVLQSGFRKME (long), in between the mNeonGreen and nanoLuc proteins. Co-expression of the sensor with the M^pro^ in live cells resulted in its cleavage in a dose- and time-dependent manner while mutation of the critical C145 residue (C145A) in M^pro^ completely abrogated the sensor cleavage. Importantly, the BRET-based sensors displayed increased sensitivities and specificities as compared to the recently developed FlipGFP-based M^pro^ sensor. Additionally, the sensors recapitulated the inhibition of M^pro^ by the well-characterized pharmacological agent GC376. Further, in vitro assays with the BRET-based M^pro^ sensors revealed a molecular crowding-mediated increase in the rate of M^pro^ activity and a decrease in the inhibitory potential of GC376. The sensor developed here will find direct utility in studies related to drug discovery targeting the SARS-CoV-2 M^pro^ and functional genomics application to determine the effect of sequence variation in M^pro^.

## Introduction

COVID-19 has become a global health threat with more than 300 million infections and more than 5 million deaths as of January 2022. The causative agent, Severe Acute Respiratory Syndrome Corona Virus 2 (SARS-CoV-2) of the *beta-coronavirus* family shares 79% similarity with SARS-CoV and 50% similarity with MERS-CoV (Middle East respiratory syndrome coronavirus)^1, 2^. SARS-CoV-2 infection cycle is initiated by the processing of two polypeptides, pp1a and pp1ab, bearing the non-structural proteins^3^ by the auto-catalytically released viral proteases, 3-chymotrypsin-like cysteine protease (3CL^pro^) or main protease (M^pro^)^4, 5^, and papain-like protease (PL^pro^)^6-8^. M^pro^ functions as a homodimer with each monomer containing an active site formed by a conserved catalytic dyad of Cys-His^7, 9^, and cleaves the large polyprotein pp1ab at 11 sites^7^. Specifically, M^pro^ recognizes a highly conserved core sequence with a critical Gln residue for cleavage^10-13^. Importantly, M^pro^ cleavage sequences are not known to be recognized by human proteases, thus making M^pro^ an attractive target for anti-SARS-CoV-2 therapy^14^.

Given the critical role played by M^pro^ in SARS-CoV-2 infection and the cleavage specificity, a number of assays have been developed to monitor the proteolytic activity of M^pro^. Genetic reporter assays based on fluorescence and bioluminescence provide sensitive and effective systems to assess the cellular functions including cell signaling, protein dimerization, conformational changes of proteins and protein-protein interactions in live cells^15-18^. Researchers have developed fluorescence and bioluminescence-based reporter assays for screening antiviral molecules against various coronaviruses (e.g. FRET^19^, split-luciferase^20, 21^ assays). Specifically, a number of studies have utilized fluorescence resonance energy transfer (FRET)-based in vitro assays, wherein peptide substrates containing the M^pro^ cleavage sequences are used as reporter, for the identification of antivirals against SARS-CoV-2 M^pro22-26^. Additionally, a FRET-based assay was utilized for the identification of Boceprevir, GC376, and calpain inhibitors II, XII as potent inhibitors of SARS-CoV-2 M^pro27^. On the other hand, a FlipGFP-based construct containing the M^pro^ N-terminal autocleavage site has been developed to screen the antivirals against SARS-CoV-2^28, 29^. In such a construct, M^pro^-mediated cleavage of FlipGFP in live cells results in the generation of the fluorescent form of GFP from the non-fluorescent form. A later report by Drayman *et al*.,have combined FlipGFP and luciferase assays to successfully identify additional M^pro^ inhibitors^30^.

In addition to the above, Bioluminescence Resonance Energy Transfer (BRET) has been used in developing a range of genetically encoded, live cell sensors^31-34^. BRET relies on the non-radiative resonance energy transfer from a light emitting luciferase protein (donor) upon oxidation of its substrate to a fluorescent protein (acceptor) with an excitation spectrum overlapping with the luciferase emission spectra. In addition to the spectral overlap, BRET also depends on the physical distance and relative orientation of the donor and the acceptor proteins.^17, 18, 35, 36^. The latter has been successfully utilized in generating a variety of molecular sensors including detecting small molecules^37-40^, structural changes in proteins^38,41^. While a number of donor-acceptor pairs with distinct spectral and energy transfer efficiencies have been utilized for BRET-based sensor development^42^, the combination of mNeonGreen (mNG)^43-46^, a bright green fluorescent protein, and nanoLuc (NLuc)^47, 48^, a small bright and stable luciferase have gained significant usage in the recent times including proteolytic cleavage sensors^**39**,**49**^ due to excellent spectral overlap and light emission characteristics^50-56^.

In the present study, we have engineered BRET-based M^pro^ proteolytic activity sensors by inserting the M^pro^ N-terminal autocleavage sequences (either the short AVLQSGFR^4, 26^ or the long KTSAVLQSGFRKME^5, 26^ in between the mNeonGreen (mNG; acceptor) and the nanoLuc luciferase (NLuc; donor) in a single fusion construct. The sensor constructs showed robust cleavage activity in live cells when coexpressed with the wild type M^pro^, both in a dose-dependent and time-dependent manner, but not in the presence of the catalytically dead C145A mutant M^pro57-59^, and with a faster kinetics and higher specificity compared to the FlipGFP-based M^pro^ sensor. We have further determined the utility of the sensors in pharmacological inhibition of the M^pro^ using the well-established M^pro^ inhibitor, GC376^27, 60-62^.Additionally, in vitro assays showed a molecular crowding-mediated proteolytic cleavage rate enhancement and inhibitor potency decrease.

## Materials & Methods

### M^pro^ N-terminal autocleavage sequence analysis

A total of 1984 sequences for the SARS-CoV-2 pp1a polyprotein available at the NCBI Virus database (https://www.ncbi.nlm.nih.gov/genome/viruses/) were downloaded and aligned using MAFFT server (https://mafft.cbrc.jp/alignment/server/)^63, 64^. The aligned sequences of the pp1a polyprotein were analyzed for the conservation of the M^pro^ N-terminal autocleavage positions (AVLQSGFR) (Fig. 1).

**Figure 1.**
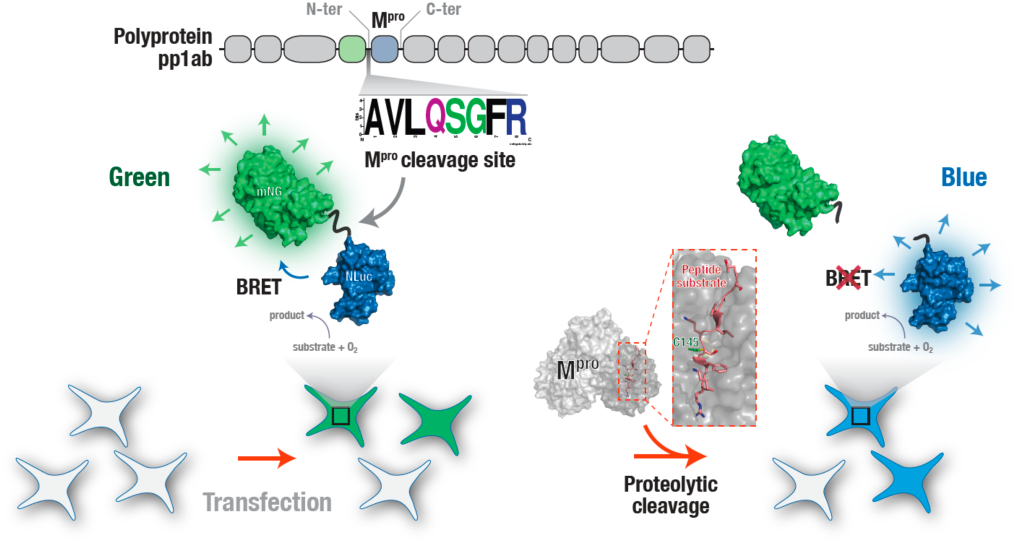
Genetically encoded, BRET-based live cell SARS-CoV-2 M^pro^ protease activity sensor. A schematic representation of the genetically encoded, BRET-based SARS-CoV-2 M^pro^ protease activity sensor expressed in live cells. Close positioning of the NLuc and mNG proteins result in a significant resonance energy transfer in the absence of the SARS-CoV-2 M^pro^ protease activity. Activity of the SARS-CoV-2 M^pro^ results in the cleavage of the sensor resulting in a decrease in the resonance energy transfer between NLuc and mNG leading to a decrease in the green fluorescence of the sensor. Shown in grey is the surface representation of M^pro^ structure (SARS-CoV; PDB: 2Q6G^65^) highlighting the critical C145 residue (green) required for proteolytic activity and the peptide substrate peptide (TSAVLQSGFRK; red).

*Structural modeling of M*^*pro*^ *N-terminal autocleavage peptide sequences* The crystal structure of the N-terminal peptide substrate complexed with SARS-CoV main protease H41A mutant (PDB: 2Q6G^65^, Chain D, aa seq: TSAVLQSGFRK) was used as a template for generating the 3D models for the short and long M^pro^ cleavage peptides, including the linker region, of the Mpro sensor (short cleavage peptide aa seq: EFGTENLYAVLQSGFRGSGGS, long cleavage peptide aa seq: EFGTENLYKTSAVLQSGFRKMEGSGGS). Models were generated using MODELLER (10.1 release, Mar. 18, 2021)^66^. Briefly, the short and long sequences were aligned with the template in PIR format. For each peptide, 100 models were initially generated using “Automodel” function and “very-slow” MD refining mode.Scoring functions such as modpdf, DOPE, and GA34, were used to assess the generated models. The model with the lowest DOPE score was further refined by loop modelling using very-slow loop MD refining mode to generate 100 refined models. The same scoring functions were used to assess the refined models. The stereochemical quality of the final model was assessed with PROCHECK ^67^.

### Molecular dynamics simulation

To neutralize the positive and negative charges on the peptide ‘s termini, the N- and C-termini were capped with N-acetyl and N-methyl amide capping groups, respectively. Topology and parameter files were generated using CHARMM-GUI webserver^68^. The biomolecular simulation systems included the peptide model, with all hydrogens added, solvated in TIP3P (transferable intermolecular potential with 3 points)^69^ cubic water box with 10 Å minimum distance between edge of box and any of the peptide atoms. Charges were neutralized by adding 0.15 M NaCl to the solvated system. The total number of atoms was 15480 and 18233 for the short and long peptide simulation systems, respectively. In silico molecular dynamics simulations were performed using Nanoscale Molecular Dynamics (NAMD) software^70^ version 2.13 with the CHARMM36(m) force field^71^. A 2 fs time-step of integration was set for all simulations performed. First, energy minimization was performed on each system for 1000 steps (2 ps). Following energy minimization, the system was slowly heated from 60 K to 310 K at 1 K interval to reach the 310 K equilibrium temperature using a temperature ramp that runs 500 steps after each temperature increment. Following thermalization, temperature was maintained at 310 K using Langevin temperature control and at 1.0 atm using Nose-Hoover Langevin piston pressure control^72^. The system was then equilibrated with 500000 steps (1 ns) using Periodic Boundary Conditions. The NAMD output structure was then used as an input for Gaussian accelerated molecular dynamics (GaMD) simulation utilizing the integrated GaMD module in NAMD and its default parameters^73, 74^ which included 2 ns of conventional molecular dynamics (cMD) equilibration run in GaMD, to collect potential statistics required for calculating the GaMD acceleration parameters, and another 50 ns equilibration run in GaMD after adding the boost potential^74, 75^, and finally GaMD production runs for 1000 ns. Both equilibration steps in GaMD were preceded by 0.4 ns preparatory runs. All GaMD simulations were run at the “dual-boost” level by setting the reference energy to the lower bound, i.e., E = Vmax^76^. One boost potential is applied to the dihedral energetic term and the other to the total potential energetic term. The details for calculating the boost potentials including the equations used have been described previously^73, 76, 77^. The upper limits of standard deviation (SD) of the dihedral and total potential boosts in GaMD were set to 6.0 kcal/mol. All GaMD simulations were performed using similar and constant temperature and pressure parameters. For all simulations, short-range non-bonded interactions were defined at 12 Å cut-off with 10 A switching distance, while Particle-mesh Ewald (PME) scheme was used to handle long-range electrostatic interactions at 1 Å PME grid spacing. Trajectory frames were saved every 10,000 steps (20 ps) and trajectory analysis was performed using the available tools in Visual Molecular Dynamics (VMD) software^78^. Trajectory movies were compiled based on 1000 frames using Videomach (http://gromada.com/videomach/) to generate 41 s movies (24 fps) in AVI format.

### M^pro^ BRET sensor plasmid construct generation

The BRET-based M^pro^ activity sensors were developed based on previously reported M^pro^ N-terminal autocleavage peptides, namely AVLQSGFR^4^ (nucleotide sequence 5 ‘ GCA GTG CTC CAA AGC GGA TTT CGC 3 ‘) and KTSAVLQSGFRKME^5, 26^(nucleotide sequence 5 ‘ AAA ACG AGT GCC GTA TTG CAG AGT GGG TTT CGG AAA ATG GAA 3 ‘), referred to as mNG-M^pro^-Nter-auto-NLuc and mNG-M^pro^-Nter-auto-L-NLuc, respectively. For this, fragments BstXI-mNG-M^pro^-Nter-auto-NLuc-XhoI and BstXI-mNG-M^pro^-Nter-auto-L-NLuc-XhoI were synthesized (Integrated DNA Technologies, IDT; Iowa, USA) and inserted into pIDTSmart (Kan) vectors to generate the plasmid constructs pIDT-mNG-M^pro^-Nter-auto-NLuc and pIDT-mNG-M^pro^-Nter-auto-L-NLuc, respectively. Both vectors were transformed into *E. coli* for amplification and purified using Qiagen mini-prep kit. Restriction enzymes BstX-I and XhoI were used to excise the two DNA fragments of interests from entry clones pIDT-mNG-M^pro^-Nter-auto-NLuc and pIDT-mNG-M^pro^-Nter-auto-L-NLuc and ligated into similarly digested destination plasmid pmNeonGreen-DEVD-NLuc [Addgene: 98287]^49^ and further confirmed by Sanger sequencing. One Shot TOP10 Competent *E. coli* cells were transformed with 2 μL of the ligation reaction and plated in LB agar plates with 100 μg/ml ampicillin. Sequences were confirmed by Sanger sequencing^79^ using a pair of forward and reverse primers, 5 ‘ GCACAGCCAGAACCACATATACCTT 3 ‘and 5 ‘ CACCACCTTGAAGATCTTCTCGATCT 3 ‘, respectively. For bacterial expression and purification of the M^pro^ sensor, the mNG-M^pro^-Nter-auto-NLuc plasmid construct was digested with *Hind*III and *Xho*I and the mNG-M^pro^-Nter-auto-NLuc fragment was subcloned into similarly digested pET-28b(+) plasmid.

### Cell culture and transfection

All experiments reported in the manuscript were performed with HEK 293T cells, which were grown in Dulbecco ‘s Modified Eagle Medium (DMEM) supplemented with 10% fetal bovine serum, and 1% penicillin-streptomycin and grown at 37 °C in 5% CO2^18, 38, 39, 41, 80-84^. Transfections were performed with polyethyleneimine (PEI) lipid according to manufacturers ‘ protocol. Briefly, HEK 293T cells were seeded onto 96-well white plates before 24 h of transfection. The plasmid DNA (sensor and M^pro^), Opti-MEM (Invitrogen; 31985088) and 1.25 μg/well of PEI lipid (Sigma-Aldrich; 408727-100mL) were combined using pipetting and incubated at room temperature for 30 min before being added to cells by droplet. The PEI stock solution of 2 mg/mL was prepared by diluting in sterile Milli-Q water and stored at -80 °C.

### Live cell, BRET-based M^pro^ proteolytic cleavage activity assays

Live cell M^pro^ proteolytic cleavage activity assays were performed by co-transfecting HEK 293T cells with either the pmNG-M^pro^-Nter-auto-NLuc or the pmNG-M^pro^-Nter-auto-L-NLuc M^pro^ sensor plasmid constructs along with either pLVX-EF1alpha-SARS-CoV-2-nsp5-2xStrep-IRES-Puro (M^pro^ WT) (a gift from Nevan Krogan (Addgene plasmid # 141370; http://n2t.net/addgene:141370 ; RRID:Addgene_141370)^85^or pLVX-EF1alpha-SARS-CoV-2-nsp5-C145A-2xStrep-IRES-Puro (C145A mutant M^pro^) plasmid (a gift from Nevan Krogan (Addgene plasmid # 141371 ; http://n2t.net/addgene:141371 ; RRID:Addgene_141371)^85^ in 96-well white flat bottom plates (Nunc; 136101). For dose-response experiments, the filler plasmid (a pcDNA3.1-based plasmid) is also co-transfected. In case of time-course experiment, a pcDNA3.1-based plasmid was used as a control (No M^pro^). The time-course experiments were carried out at 1:5 reporter-to-protease ratio. Post 48 h (or otherwise indicated) of transfection, BRET measurements were performed by the addition of furimazine (Promega, Wisconsin, USA) at a dilution of 1:200. In time-course experiments, BRET was measured at the indicated time points. Experiments were performed in triplicates and repeated a minimum of two times.

### Western blot analysis

HEK 293T cells co-transfected with the M^pro^ sensor and the M^pro^ (wild-type or mutant) plasmids were lysed in 200 μl of 2x Laemmli sample buffer (50 mM Tris-Cl pH 6.8, 1.6% SDS, 8% glycerol, 4% β-mercaptoethanol and 0.04% bromophenol blue) (heated to 85°C and sonicated prior to addition). Equal volumes of the cell lysates (30 mL) were separated by 10% SDS-PAGE using running buffer (25 mM Tris, 192 mM glycine, 0.1% SDS) at a constant voltage of 100 V for 1.5 h following which proteins were transferred onto PVDF (Polyvinylidene fluoride) membranes. Membranes were blocked in Tris-buffered saline containing 0.1% Tween-20 (TBS-T) with skimmed milk (5%) for 1 h at room temperature. Blots were incubated either with anti-His antibody (6x-His Tag Monoclonal Antibody (HIS.H8), Alexa Fluor 488; ThermoFisher Scientific-MA1-21315-A488; 1:5000) or with anti-Strep-tag mouse monoclonal antibody (anti-Strep-tag mouse monoclonal, C23.21; PROGEN-910STR; 1:5000) overnight at 4°C in dilution buffer (TBS-T containing 5% bovine serum albumin (BSA). Secondary anti-mouse IgG HRP (Anti-Mouse Ig:HRP Donkey pAb; ECM biosciences-MS3001; 1:10000 diluted in TBS-T) was used to detect M^pro^ and the cleaved M^pro^ sensor proteins.

### Live cell M^pro^ proteolytic cleavage inhibitor assay

HEK 293T cells were co-transfected with either pmNG-M^pro^-Nter-auto-NLuc or pmNG-M^pro^-Nter-auto-L-NLuc plasmid along with either pLVX-EF1alpha-SARS-CoV-2-nsp5-2xStrep-IRES-Puro (M^pro^ WT) (Addgene plasmid # 141370) or pLVX-EF1alpha-SARS-CoV-2-nsp5-C145A-2xStrep-IRES-Puro (M^pro^ C145A) (Addgene plasmid # 141371) plasmid in 96-well white flat bottom plates. We have used 1:5 reporter-to-protease ratio for the assay. Eight hours post-transfection, GC376 (GC376 Sodium; AOBIOUS-AOB36447; stock solution prepared in 50% DMSO at a concentration of 10 mM) was added to the cells at different concentrations. After 24 h of incubation with the inhibitor, BRET measurements were performed by the addition of furimazine (Promega, Wisconsin, USA) at a dilution of 1:200. The percentage activity was calculated by normalizing the BRET ratio with the negative control (No M^pro^). Two independent experiments were performed in triplicates for each sensor construct.

### Live cell, FlipGFP-based M^pro^ proteolytic assay

For live cell FlipGFP-based M^pro^ proteolytic activity assays, HEK 293T cells were seeded onto 24-well plates and co-transfected with the FlipGFP sensor plasmid (pcDNA3 FlipGFP(Mpro) T2A mCherry; a gift from Xiaokun Shu; Addgene plasmid # 163078)^29^ and either the WT or the C145A mutant M^pro^ expressing plasmid DNA (1.25 μg/well) using PEI lipid after 24 h of cell seeding. For transfection, cells were imaged using a EVOS FL microscope (Life Technologies; 4’ objective) at the indicated time in the red (to monitor mCherry expression to determine transfected cells) and the green (to monitor conversion of non-fluorescent FlipGFP into the fluorescent GFP form after M^pro^-mediated cleavage) channels. Images were analyzed for percentage GPF positive (GPF^+^) cells, number of transfected cells and total number of analyzed cells for each time point using Fiji^86^. The ImageJ macro script used for the analysis is provided in the Supporting Text. For determining number of GFP^+^ cells, GFP intensities obtained for each cell was background corrected and threshold was applied.

### Cell lysate preparation for in vitro BRET assays

To prepare cell lysates containing the M^pro^ sensors, HEK 293T cells were transfected with either the mNG-M^pro^-Nter-auto-NLuc or the mNG-M^pro^-Nter-auto-L-NLuc M^pro^ BRET sensor and washed with chilled Dulbecco ‘s Phosphate-Buffered Saline (DBPS) 48 h post transfection. Cells were lysed in a buffer containing 50 mM HEPES (pH 7.5), 50 mM NaCl, 0.1% Triton-X 100, 1 mM Dithiothreitol (DTT) & 1 mM ethylenediamine tetraacetic acid (EDTA)^87^ on ice. Cell lysates were collected in a 1.5 mL Eppendorf tube and centrifuged at 4 °C for 1 h at 14,000 rotations per min (RPM) following which supernatant were collected and stored at -80 °C until further usage.

### In vitro, BRET-based M^pro^ proteolytic cleavage activity assays

In vitro BRET-based M^pro^ proteolytic cleavage activity assays were performed by incubating cell lysates containing the short, BRET-based M^pro^ sensor with different concentrations (0.5, 5, 50 and 500 nM) of recombinantly purified SARS-CoV M^pro^ (SARS coronavirus, 3CL Protease, Recombinant from E. coli; NR-700; BEI Resources, NIAID, NIH; stock solution of the protein was prepared by dissolving the lyophilized protein in 50 μM in Tris-buffered saline (TBS) containing 10% glycerol) and BRET monitored through luminescence scans. The effect of molecular crowding was monitored by incubating the sensor and the protease in the absence or presence of 25% (w/v) of polyethylene glycol (PEG) of a range (0.4, 2, 4, 8, 20 or 35 kDa) of molecular weights (Sigma-Aldrich). GC376 (GC376 Sodium; AOBIOUS -AOB36447; stock solution prepared in 50% DMSO at a concentration of 10 mM) inhibition of M^pro^ (50 nM) activity was monitored under a range of the inhibitor concentrations in the absence or presence of 25% (w/v) PEG 8K. BRET measurements were performed at 37°C by the addition of furimazine (Promega, Wisconsin, USA) at a dilution of 1:200. The bioluminescence (467 nm) and fluorescence (533 nm) readings were recorded using Tecan SPARK multimode microplate reader and used to calculate the BRET ratios (533 nm/467 nm). Total mNG fluorescence in cell lysates containing the short, BRET-based M^pro^ sensor was measured by exciting the samples at 480 nm and emission acquired at a wavelength of 530 nm.

### BRET and fluorescence measurements

BRET measurements were performed using a Tecan SPARK® multimode microplate reader. Bioluminescence spectral scan was performed from 380 to 664 nm wavelengths with an acquisition time of 400 ms for each wavelength to determine relative emissions from NLuc (donor) and mNG (acceptor) and quantify BRET, which is expressed as a ratio of emissions at 533 nm and 467 nm. In some experiments, BRET measurements were performed by measuring emission only at 533 and 467 nm. Total mNG fluorescence in the sensor expressing cells was measured by exciting the samples at 480 nm and emission acquired at a wavelength of 530 nm.

### Data analysis and figure preparation

GraphPad Prism (version 9 for macOS, GraphPad Software, La Jolla California USA; www.graphpad.com), in combination with Microsoft Excel, was used for data analysis and graph preparation. Figures were assembled using Adobe Illustrator.

## Results and Discussion

### BRET-based M^pro^ proteolytic cleavage activity sensor design

In order to develop a live cell, BRET-based specific reporter to monitor M^pro^ proteolytic cleavage activity, we generated fusion proteins containing the M^pro^ N-terminal autocleavage sequence sandwiched between mNG (acceptor) and NLuc (donor) proteins (Fig. 1). The mNG and NLuc pair (acceptor and donor, respectively) has been used in a number of BRET-based sensor and show efficient energy transfer from NLuc to mNG. Thus, in the absence of any proteolytic cleavage, the sensor constructs are expected to display significant emission in the green channel. However, upon proteolytic cleavage of the sandwiched autocleavage peptide, the sensor constructs will display reduced emission in the green channel with a concomitant increase in the emission in the blue channel (Fig. 1).

Both SARS-CoV-2^10^ and SARS-CoV-1^87^ M^pro^ show a significant preference for the N-terminal autocleavage sequence (AVLQSGFR; short sensor; Fig. 1,2A; see Supporting Text for complete sequence of the sensor constructs) as a substrate compared to other cleavage sequences in the pp1a polyprotein in terms of catalytic efficiency, and has been widely utilized in FRET-based, in vitro assays^22, 26, 28, 88^ as well as in a FlipGFP-based, live cell assay^28^. Additionally, we analyzed all available pp1a polyprotein sequences reported for SARS-CoV-2 isolates at the NCBI Virus database for any variation in the cleavage sequence. This indicated that the N-terminal autocleavage sequence is invariable in all isolates reported and therefore, the sensor construct designed here will serve as a reliable reporter for M^pro^ proteolytic cleavage activity.

**Figure 2.**
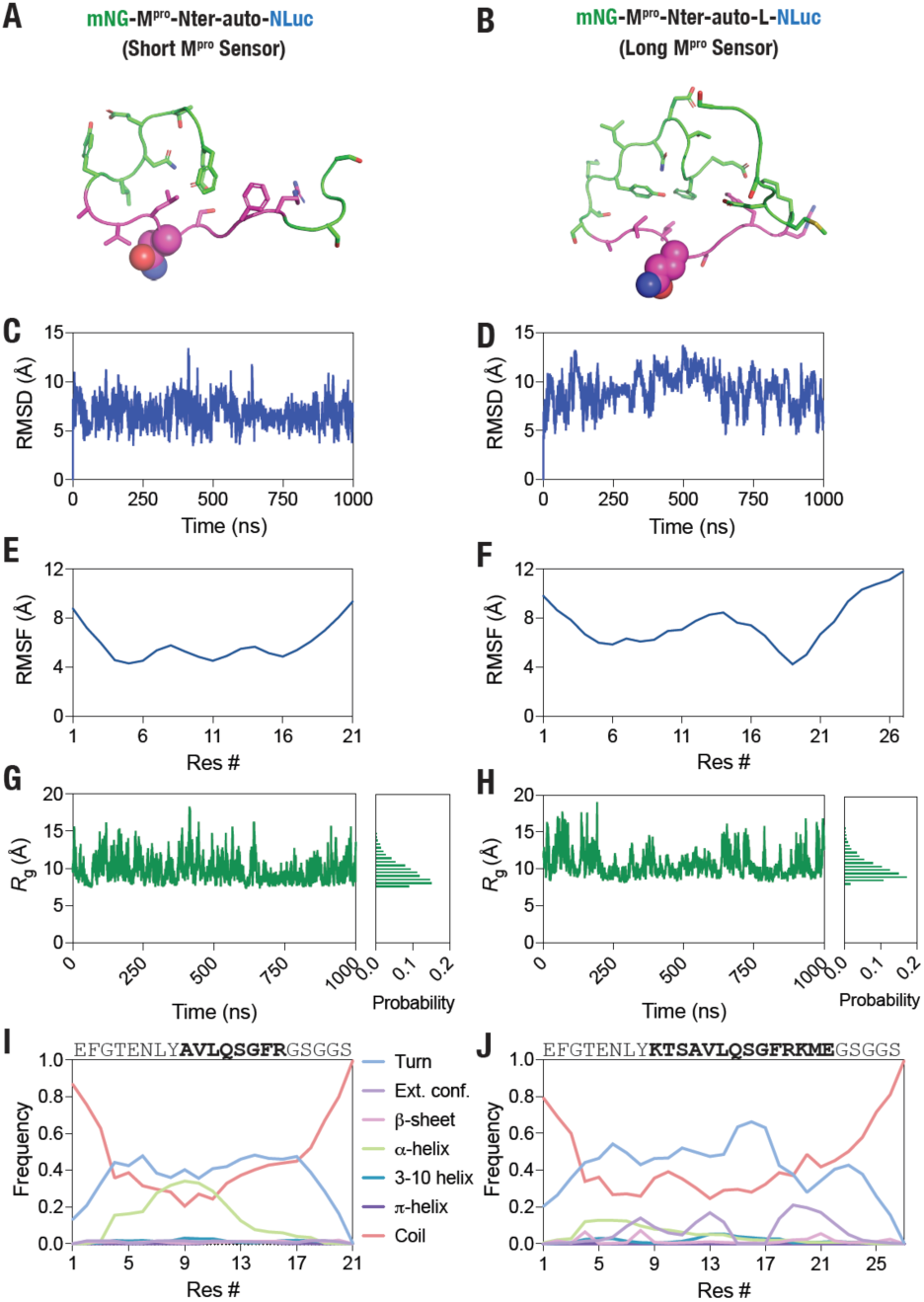
M^pro^ N-terminal autocleavage peptide is flexible. (A,B) Cartoon representation of the M^pro^-Nter-auto (short; A) and M^pro^-Nter-auto-L (long; B) peptide structures modeled using the peptide substrate crystallized with H41A mutant SARS-CoV M^pro^ (PDB: 2Q6G^65^). (C,D) Graph showing backbone (Cα) root-mean-square deviation (RMSD) values of M^pro^-Nter-auto (short; C) and M^pro^-Nter-auto-L (long; D) peptide obtained from 1 ms of GaMD simulations. (E,F) Graph showing backbone (Cα) root-mean-square fluctuation (RMSF) values of M^pro^-Nter-auto (short; E) and M^pro^-Nter-auto-L (long; F) peptides. (G,H) Graph showing radius of gyration (Rg) of the M^pro^-Nter-auto (short; G) and M^pro^-Nter-auto-L (long; H) peptides monitored over 1 ms of GaMD simulations. (I,J) Graph showing frequency of indicated secondary structures formed by the M^pro^-Nter-auto (short; I) and M^pro^-Nter-auto-L (long; J) peptides over the 1 ms of GaMD simulations. T, turn; E, extended conformation; B, isolated bridge; H, α-helix; G, 3-10 helix; I, π-helix; C, coil;

We note that while BRET comes with several advantages including a higher signal-to-noise ratio and an extended dynamic range compared to some other methods^89^, the presence of the acceptor and donor proteins i.e. mNG and NLuc at the N- and C-termini, respectively could potentially affect the interaction of the cleavage peptide with the M^pro^ dimer and thus, in turn, affect the cleavage efficiency of the peptide. This is especially relevant given that the binding of the peptide substrate has been reported to allosterically activate the SARS-CoV-1 M^pro^ dimer. Therefore, we generated a second, extended M^pro^ sensor construct the KTSAVLQSGFRKME peptide sequence (containing additional three residues on each sides of the AVLQSGFR core sequence; long sensor; Fig.3B; see Supporting Text for complete sequence of the sensor construct), which has also been used in several previous reports^5, 22, 26^. A key requirement for efficient cleavage of peptide substrates by M^pro^ is their structural flexibility as previous studies have reported the formation of secondary structural element can alter cleavage activity^90-92^, especially given that secondary structure prediction indicated a-helical propensity by both the short as well as the long peptide (Supp. Fig. 1). In order to assess structural flexibility and secondary structure formation by the two peptides, we generated structural models of the peptides using the substrate peptide co-crystalized with the H41A mutant SARS-CoV-1 M^pro65^ and performed all-atom, explicit solvent, GaMD simulations that allow enhanced sampling of protein conformational states by applying harmonic boost potential that follows Gaussian distribution to accelerate conformational transitions^73, 74^. Structural models were generated using Modeler^66^ (Fig. 2A,B) and MD simulations were performed using the NAMD software^93^ for a total duration of 1 ms for each peptide (Supp. Movies 1,2). These simulations indicated significant structural fluctuations in the two peptides as revealed by relatively large root-mean-squared-deviation (RMSD) and root-mean-squared-fluctuations (RMSF) (Fig. 2C,D). Further, radius of gyration (*R*_g_) measurements of the peptides over the course of simulation also revealed structural fluctuations of the peptides with an appreciably greater fluctuations observed for the short peptide compared to the long one (Fig. 2G,H). Additionally, this analysis also revealed a greater *R*_g_ for the longer peptide compared to the short peptide (median *R*_g_ values of 9.4 vs. 10.2 Å) as expected. Finally, secondary structure analysis of the peptides over the course of the 1 ms long simulation trajectory revealed that the peptides largely show a propensity to form turns (Fig. 2I, J). Notably, certain central residues in the shorter peptide may form a-helix that was not seen with the longer peptide leading to the possibility of a differential cleavage efficiency of the peptides by M^pro^. These results were in agreement with dynamic light scattering (DLS) measurements that showed an average size of 6.2 ± 1.2 nm for the short M^pro^ biosensor (Fig. S1C), suggesting a compact structure of the sensor construct, which may show high BRET signal that is critical for detecting sensor cleavage. In the following, we report experimental results with both the sensor constructs in order to provide a comparative analysis and determine the one that serves as a better substrate and thus, provide a superior evaluation of M^pro^ proteolytic cleavage activity in live cells.

**Figure 3.**
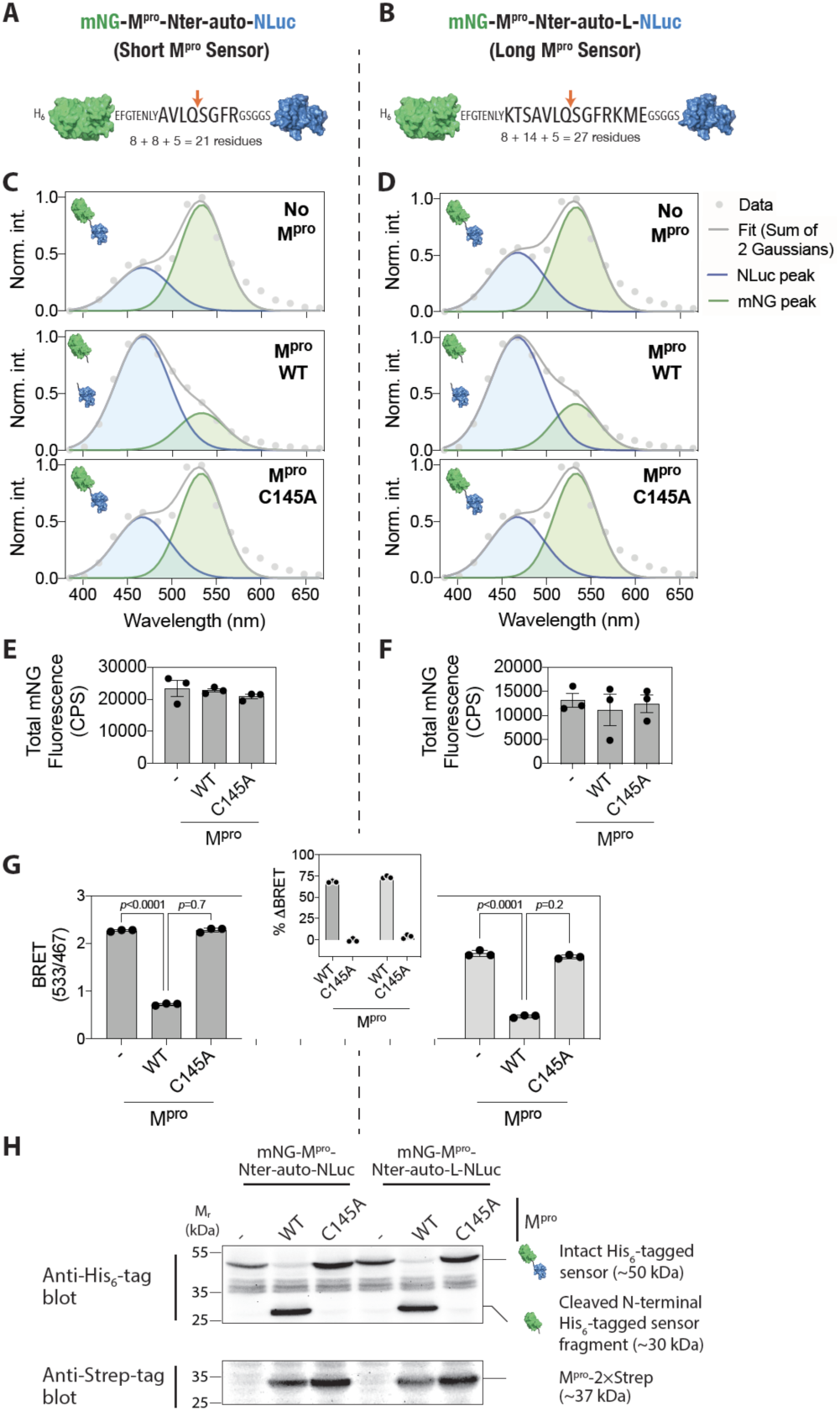
Efficient cleavage of the M^pro^ sensor constructs in live cells. (A,B) Schematic showing the M^pro^ sensor constructs -short (A) and long (B) – with SARS-CoV-2 M^pro^ N-terminal autocleavage sequence. (C,D) Graph showing a representative bioluminescence spectra of the short (C) and long (D) M^pro^ sensor constructs either in control cells or in cells expressing the WT or C145A mutant M^pro^. Data were fit to a two Gaussian model reflecting mNG fluorescence and NLuc bioluminescence peaks. Note the reduction in the mNG peak (533 nm) of both the short and the long sensors when coexpressed with the wild type M^pro^ while no reduction was observed when coexpressed with the C145A mutant M^pro^. (E,F) Graphs showing total mNG fluorescence (measured prior to substrate addition) in cells expressing the short (E) and the long (F) sensors. (G) Graph showing BRET ratio (ratio emission at 533 nm and 467 nm) of the short (left side) and the long (right side) M^pro^ protease activity sensors in either control cells or when coexpressed with the wild type or the C145A mutant M^pro^. Inset: graph showing percentage change in BRET of the short (left side) and the long (right side) when co-expressed with the wild type or the C145A mutant M^pro^. Data shown are mean ± S.D. from a representative of independent experiments performed multiple times. (H) Top panel: anti-His tag blot showing cleavage of the short (left side) and the long (right side) M^pro^ sensor constructs in either control cells or in cells co-expressing the wild type or the C145A mutant M^pro^. Note the release of an approximately 30 kDa, His_6_-tagged-mNG fragment in cells expressing the wild type but not in the C145A mutant M^pro^.Bottom panel: anti-Strep-tag blot showing expression of the M^pro^ in the respectively transfected cells.

### BRET-based M^pro^ proteolytic cleavage activity sensor characterization in live cells

In order to test the functionality of the BRET-based M^pro^ proteolytic cleavage activity sensors, we transfected HEK 293T cells with the short and long sensors either alone or along with the M^pro^ expressing plasmid in a 1:5 sensor-to-protease plasmid ratio (Fig. 3). Additionally, we utilized the catalytic dead C145A mutant M^pro^ as a negative control in these experiments since Cys145 is essential for the proteolytic activity of M^pro57-59^. The transfection efficiency and expression of the sensor constructs was monitored by imaging live cells for mNG fluorescence using an epifluorescence microscope, which showed an efficient transfection and expression of the sensor constructs after 24 h of transfection (Supp. Fig. S2).

We then determined the spectral properties of the two sensors in live cells. For this, sensor construct transfected cells in adherent conditions were incubated with NLuc substrate and emission in the range of 380 and 664 nm wavelength were detected using a microplate reader. In the absence of coexpression of M^pro^, both the short and the long sensors showed two peaks corresponding to NLuc (467 nm) and mNG (533 nm), respectively, as determined from two Gaussian fitting of the spectral data (Fig. 3C,D; top panels). Coexpression of the wild type M^pro^ resulted in a decrease in the mNG emission peak in cells expressing either of the sensor constructs (Fig. 3C, D; middle panel) while no such decrease was observed when the C145A mutant M^pro^ was coexpressed with the sensor constructs (Fig. 3C,D; bottom panel). We note that the coexpression of either the wild type or the C145A mutant M^pro 57-59^ did not result in any significant change in the intracellular levels of the sensor constructs as determined from mNG fluorescence at 530 nm under excitation with 480 nm light (Fig. 3E,F).

We then determined the BRET ratio of the sensor constructs in live cells under different M^pro^ coexpression conditions as a ratio of emission at 533 nm and 467 nm. Basal BRET ratio of the short and long sensors were found to be 2.37 ± 0.17 vs 1.79 ± 0.06 (mean ± standard deviation; *N* = 6 each; independent experiments performed in triplicates; *p* < 0.0001), respectively, indicating that the additional 6 residues in the long sensor resulted in a 24 (± 2)% decrease in the BRET ratio. Coexpression of the wild type M^pro^ resulted in a significant decrease in the BRET ratio while no significant decrease was observed in the presence of the C145A mutant M^pro^ (Fig. 3G).Importantly, both the short and the long sensor expressing cells showed ∼75% reduction in the BRET ratio in the presence of the wild type M^pro^ (Fig. 3G, inset), indicating that these sensors will provide a wide dynamic range for monitoring M^pro^ proteolytic cleavage activity in live cells. Importantly, no change in the BRET ratio in cells expressing either of the sensors was observed in presence of the C145A mutant M^pro^ indicating high specificity of the BRET signals of these sensors (Fig. 3G, inset).

In order to confirm that the reductions in the BRET observed upon coexpression with M^pro^, we performed western blot analysis of cell lysates prepared from the M^pro^ sensor transfected cells. For this, we utilized the N-terminal His_6_-tag in the M^pro^ sensor constructs, which will be retained in the N-terminal, mNG protein containing fragment upon proteolytic cleavage, and C-terminal 2x Strep-tag in the M^pro^ protein for detecting cleavage of the M^pro^ sensor constructs and the expression of M^pro^, respectively. Cells transfected with only the M^pro^ sensor constructs showed a band of the expected molecular weight of ∼50 kDa (as predicted from the amino acid sequence of the sensor constructs) (Fig. 3H and Supp. Fig. S3). Indeed, cells coexpressing the wild type M^pro^, as assessed from the anti-Strep-tag blot, showed a band of ∼30 kDa corresponding to the predicted molecular weight of the cleaved N-terminal fragment containing the His_6_-tag and the mNG protein with a concomitant loss of the full-length sensor constructs (Fig. 3H). However, cells coexpressing the C145A mutant M^pro^, as assessed from the anti-Strep-tag blot, did not show the cleaved sensor fragment (Fig. 3H). Agreement of these results with the BRET measurements shown above establishes that the reduction in the BRET ratio observed in the presence of the wild type M^pro^ is due to the proteolytic cleavage of the sensor constructs, and therefore, live cell BRET ratio measurements can be reliably used as a measure of M^pro^ proteolytic activity.

### M^pro^ dose-dependent cleavage of the sensor in live cells

Having established that the BRET ratio could be used to detect M^pro^ proteolytic activity of the sensor constructs, we aimed to determine the M^pro^ dose-dependent cleavage of the sensor constructs in live cells.For this, we cotransfected the cells with the 25 ng/well sensor constructs and a range of M^pro^ plasmid concentrations (0,0.0125, 0.125, 1.25, 12.5 and125 ng/well) and monitored bioluminescence spectra in adherent cells after 48 h. This revealed a M^pro^ plasmid dose-dependent shift in the bioluminescence spectra (Supp. Fig. S4) and the BRET ratio (Fig. 4A,B) of both the short and the long sensor in the presence of the wild type M^pro^ but not in the presence of the C145A mutant M^pro^. Discernable decreases in the BRET ratio could be observed at a minimum amount of 1.25 ng of M^pro^ plasmid DNA and a maximum decrease in the BRET ratio of ∼80% at the highest concentration of 125 ng for both sensor constructs (Fig. 4A,B; insets). The analysis has also showed that the EC_50_ values are 1.09 ± 0.09 ng/well and 0.91 ± 0.89 ng/well for the short and the long sensors, respectively. These data demonstrate the functional potency of M^pro^ expressed in these cells.

**Figure 4.**
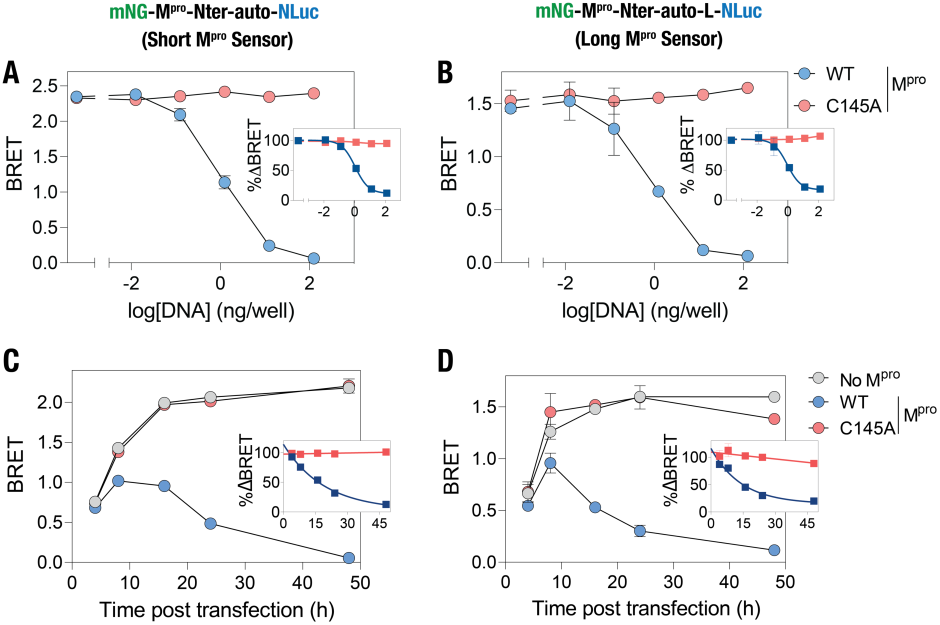
M^pro^ DNA dose- and time-dependent cleavage of the M^pro^ sensors in live cells. (A,B) Graph showing BRET ratio of the short (A) and the long (B) M^pro^ sensors in cells transfected with the indicated amounts of either the wild type or the C145A mutant M^pro^ plasmid DNA. Insets: graph showing percentage decrease in BRET ratio compared to the control cells when transfected with the indicated amounts of the wild type M^pro^ plasmid DNA. (C,D) Graphs showing the BRET ratio of the short (C) and the long (D) M^pro^ sensors at the indicated time post transfection in either control cells or cells transfected with the wild type or the C145A mutant M^pro^. Insets: graph showing percentage change in BRET ratio compared to the control cells with time when transfected with the wild type or mutant M^pro^. Data shown are mean ±S.D. from a representative of independent experiments performed thrice.

### Monitoring the temporal dynamics of M^pro^ proteolytic activity in live cells

We then monitored the temporal dynamics of M^pro^ proteolytic activity in live cells. Towards this, we transfected the cells with the M^pro^ sensor constructs either in the absence or in the presence of the wild type or the C145A mutant M^pro^ plasmid and monitored the bioluminescence spectra from 4 h until 48 h post transfection (Supp. Fig. S5). Analysis of the bioluminescence spectra obtained from cells expressing either of the M^pro^ sensors indicated a lower BRET ratio after 4 h of transfection, which increased with time and plateaued after 16 h of transfection in the absence of M^pro^ (Fig.4C,D). Although mNG shows a relatively fast maturation time compared to several other fluorescent proteins^94^, these data likely indicate a relatively slower intracellular maturation of mNG compared to NLuc^48, 95^ (Supp. Fig. 6).Importantly, a significant decrease in the BRET ratio of cells expressing either of the M^pro^ sensors could be observed in the presence of the wild type M^pro^ after 8 h of transfection (Fig. 4C,D). BRET ratio of the cells continued to decrease in the presence of the wild type M^pro^ until 48 h of transfection while no such decrease was observed in the presence of the C145A mutant M^pro^ (Fig. 4C,D; insets). The half-life of the protease was found to be 13.22 ± 3.22 h and 9.81 ± 2.54 h for the short and the long sensors, respectively. These data suggests that the BRET-based M^pro^ sensor developed here could report M^pro^ proteolytic activity as early as 8 h of infection, although this may vary depending on the actual expression of the protease in host cells.

### Comparison with the FlipGFP-based M^pro^ proteolytic sensor in live cells

Having established the monitoring of expression-dependent proteolytic activity of M^pro^ in live cells, we then performed similar experiments with the recently reported FlipGFP-based M^pro^ proteolytic activity reporter^29^to compare their performance of the biosensors in reporting M^pro^ proteolytic activity in live cells. For this, we transfected HEK 293T cells with the FlipGFP M^pro^ sensor expression plasmid along with either the WT or C145A M^pro^ expression plasmid and monitored GFP expression in the cells to ascertain conversion of the non-fluorescent protein to a fluorescent one while mCherry expression in the cells was used for detecting transfected cells. Epifluorescence imaging of the cells post 4 h of transfection revealed the appearance of GFP fluorescence in the transfected cells, as ascertained from mCherry fluorescence, in the presence of WT M^pro^ after 24 h of transfection (48 ± 2%) while more cells showed GFP fluorescence after 48 h of transfection (70 ± 4%) (Fig. 5A,B). These data indicate a delayed response of the FlipGFP sensor to M^pro^ proteolytic activity in comparison to the BRET-based sensor. Additionally, a significant number of cells were found to be GFP positive after 24 h (9 ± 1%) and 48 h (20 ± 1%) of FlipGFP transfection in the presence of the C145A mutant M^pro^ (Fig. 5C,D). This is contrast to the observations made with the BRET-based sensor in the presence of the mutant M^pro^ (Fig. 4C,D).

**Figure 5.**
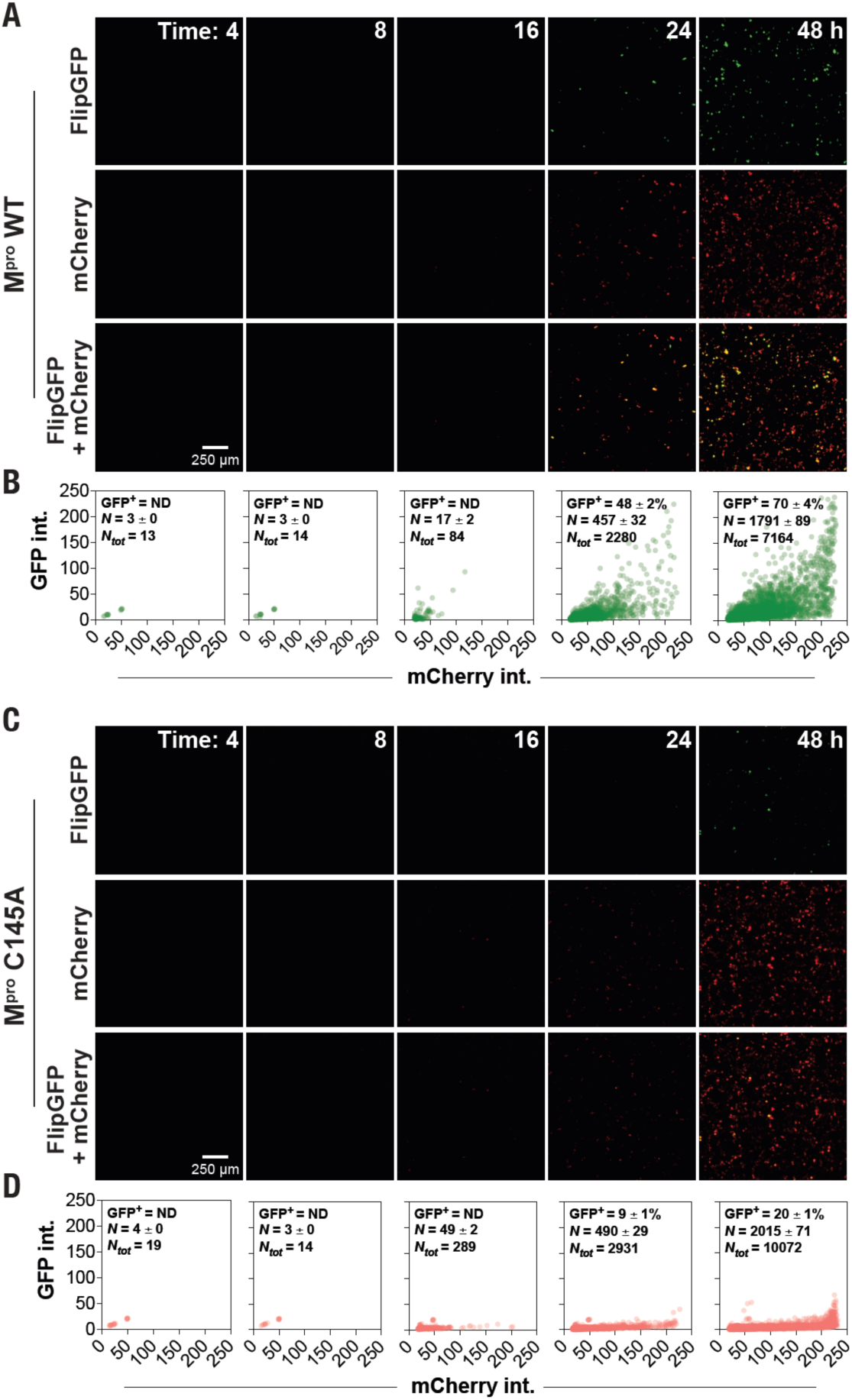
Monitoring M^pro^ proteolytic activity using the FlipGFP-based M^pro^ sensor in live cells. (A) Epifluorescence images of cells showing time-dependent expression of GFP, which is converted from the non-fluorescent FlipGFP upon proteolytic cleavage by M^pro^ (top panel), mCherry (middle panel) and merge (bottom panel) in cells transfected with the WT M^pro^.(B) Graphs showing GFP and mCherry fluorescence in individual cells transfected with the WT M^pro^ at the indicated time points. (C) Epifluorescence images of cells showing time-dependent expression of GPF (top panel), mCherry (middle panel) and merge (bottom panel) in cells transfected with the C145A mutant M^pro^.(D) Graphs showing GFP and mCherry fluorescence in individual cells transfected with the C145A mutant M^pro^ at the indicated time points. Data shown are mean ±S.D. from a representative of independent experiments performed thrice.

### Monitoring pharmacological inhibition of M^pro^ proteolytic activity in live cells

Finally, we determined the utility of the BRET-based M^pro^ sensors in pharmacological inhibition of M^pro^ proteolytic activity in live cells. Towards this, we treated cells coexpressing the M^pro^ sensors and M^pro^ with various concentrations of GC376, which has been shown to inhibit M^pro^ in live cells^27, 60-62^, after 8 h of transfection based on the results reported above, and determined bioluminescence spectra of the cells after an additional 24 h (Supp. Fig. S7). A GC376 dose-dependent increase in the BRET ratio of cells coexpressing either the short or the long sensor and the wild type M^pro^ was observed, while no sensor cleavage was observed in the presence of the C145A mutant M^pro^ (Fig. 6A,B). Percentage proteolytic cleavage activity determined from the BRET ratio indicate that GC376 starts to inhibit M^pro^ at 33.3 μM concentration and continued to do so until a concentration of 333 μM (Fig. 6A,B; insets). The data has showed that the IC_50_ values for the short sensor is 127.4 ± 23.33 μM and that for long sensor is 194.7 ± 7.49 μM. The lower efficacy of GC376 observed here compared to previous reports^27, 28, 60-62,27,58^ perhaps indicates a cell type-or M^pro^ expression-dependent effect. Taken together, these data indicate that the BRET-based M^pro^ proteolytic activity sensors developed here can be utilized for screening antivirals targeted against M^pro^.

**Figure 6.**
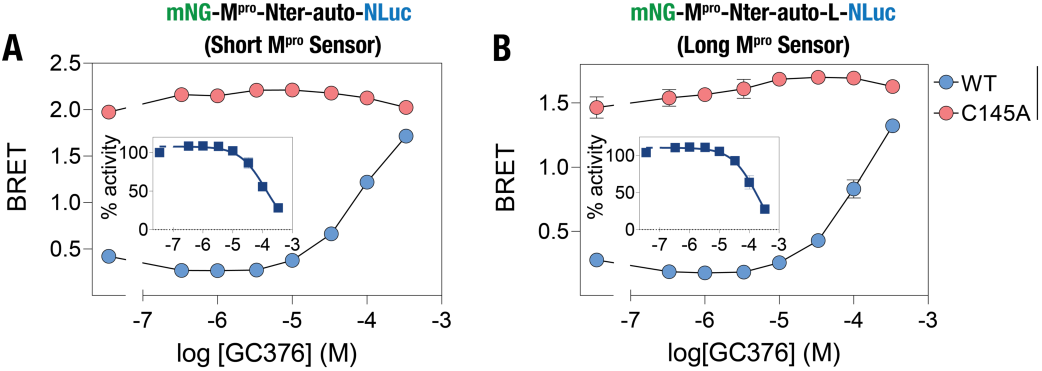
M^pro^ inhibition monitored in live cells using the BRET-based M^pro^ sensors. (A,B) Graphs showing GC376-mediated inhibition of M^pro^ proteolytic cleavage of the short (A) and long (B) M^pro^ sensor in live cells. Insets: graphs showing percentage proteolytic activity of WT M^pro^ in live cells. Data shown are mean ± S.D. from a representative of independent experiments performed thrice.

### Monitoring M^pro^ proteolytic cleavage activity in vitro

Having established the utility of the BRET-based M^pro^ sensor in live cell studies, we then focused on determining their utility in vitro using a recombinantly purified M^pro^. For this, we prepared lysates from HEK 293T expressing either the short or the long M^pro^ sensor construct, incubated equivalent amounts of the lysates with three different concentrations (5 μM, 500 nM and 50 nM) of the recombinantly purified M^pro^ and monitored BRET following addition of the NLuc substrate (Fig. 7A,B). This assay revealed a M^pro^ concentration-dependent proteolytic processing of the M^pro^ sensors as ascertained from the decreasing BRET ratios.Importantly, the assay also indicated that a minimum of 500 nM of the recombinantly purified M^pro^ is required for a discernable proteolytic cleavage of the sensors as the BRET ratio of the sensors decreased to a lesser extent in the presence of 50 nM M^pro^ protein while a substantially higher rate of cleavage was observed under 500 nM M^pro^.Enzyme kinetics assays performed using a range of the short M^pro^ sensor concentrations 200 nM of M^pro^ revealed a low *k* ‘ value of 3.09 ± 0.02 μM, which is lower than the *K*_m_ value of 17^96^ or 56^97^ μM determined using a FRET-based peptide substrate, and Hill coefficient of 1.58 (Supp. Fig. S8), suggesting a significant cooperativity in the protein. Further, an increase in M^pro^ concentration resulted in a decrease in the Hill coefficient to 1.16 (Supp. Fig. S8), suggesting a role for protein dimerization in the observed cooperativity in M^pro^.^98^ We then performed the assays in the presence of GC376 to determine the pharmacological inhibition of M^pro^ activity in vitro. For this, 500 nM M^pro^ was preincubated with a range of concentrations of GC376 (10^−4^ to 10^−9^ M) for 30 minutes at 37 °C and cleavage activity was monitored after addition of lysates prepared from cells expressing either the short or the long M^pro^ sensor. Incubation with GC376 resulted in a decrease in the rate of proteolytic cleavage of both the short and the long M^pro^ sensor (Fig. 7C,D) with IC_50_ values of 73.1 ± 7.4 and 86.9 ± 11.0 nM for the short and the long sensor, respectively (Fig.7G,H).

**Figure 7.**
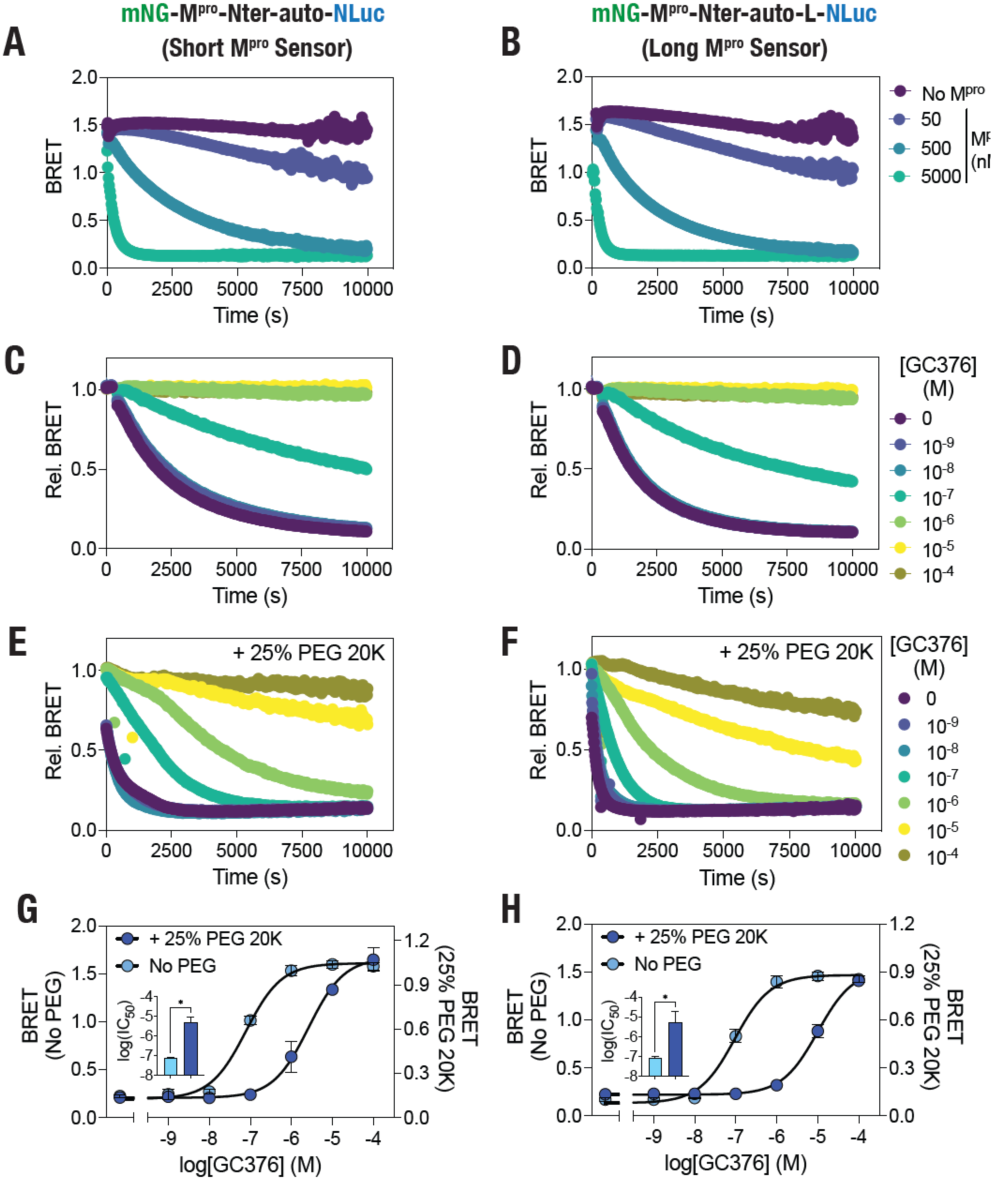
Molecular crowding-mediated increase in M^pro^ proteolytic activity and decrease in GC376 potency. (A,B) Graph showing in vitro proteolytic cleavage kinetics of the short (A) and the long (B) M^pro^ biosensor under the indicated concentrations of the recombinantly purified M^pro^ protein. (C,D) Graphs showing GC376-mediated inhibition of SARS-CoV M^pro^ proteolytic cleavage of the short (C) or the long (D) M^pro^ sensors. (E,F) Graphs showing GC376-mediated inhibition of M^pro^ proteolytic cleavage of the short (E) or the long (F) M^pro^ sensors in the presence of 25% (w/v) of PEG 20K. (G,H) Graph showing concentration-dependent inhibition of M^pro^ determined using either the short (G) or the long (H) M^pro^ sensors in the absence or in the presence of 25% (w/v) of PEG 20K. Inset in G & H: graph showing logIC_50_ values of GC376 in the absence and presence of 25% (w/v) of PEG 20K. Data shown are mean ±S.D. obtained from assays repeated thrice. (C,D) Graph showing in vitro proteolytic cleavage kinetics of the short M^pro^ biosensor in the absence and presence of 25% (w/v) of PEG 20000 (20K).

### In vitro assays reveal molecular crowding-mediated increase in M^pro^ proteolytic activity and a decrease in inhibitor efficacy

We took advantage of the slow rate of the M^pro^ sensor cleavage under 500 nM M^pro^ to determine the effect of molecular crowding on the proteolytic activity of the protein.Molecular crowding in the intracellular environment caused by the presence of soluble and insoluble macromolecules such as proteins, nucleic acids, ribosomes and carbohydrates has been shown to impact both structure and stability of proteins in cells as well as enzyme kinetics^99-104^ including a decrease in the activity HCV NS3/4A protease^105^ and an increase in the proteolytic activity of SARS-CoV M^pro103^. For this, we included 25% (weight/volume; w/v) of 20000 Da (20K) polyethylene glycol (PEG), which is a non-toxic, hydrophilic polyether that serves as a crowding agent and has been extensively utilized to simulate the molecular crowding in vitro^106, 107^, in the assays and monitored cleavage of the M^pro^ sensors under 500 nM M^pro^ under varying concentrations of GC376. Inclusion of 25% PEG 20K resulted in a substantial increase in the rate of proteolytic cleavage of the M^pro^ sensors in the absence of GC376 (Fig. 7E,F). This data suggests that molecular crowding caused by PEG 20K is likely effective in causing increased dimerization of M^pro^, a feature critical for its catalytic activity, through an increase in the effective concentration of the protein due to excluded volume effects, and thus increases the rate of proteolytic cleavage of the M^pro^ sensor. Importantly, while GC376 could inhibit M^pro^ activity, the IC_50_ values as obtained from BRET ratios after 2 hours of incubation with both the short as well as the long sensor indicated a large shift (IC_50_ values of 2623 ± 760 and 10260 ± 3280 nM, respectively) (Fig. 7G,H). Together, these data indicate that M^pro^ could be more active in the crowded environment of an infected host cell compared to in vitro conditions, and may require higher concentrations of pharmacological inhibitors for effective inhibitions of its catalytic activity than those determined from in vitro assays.

## Conclusion

To conclude, we have developed genetically encoded, BRET-based M^pro^ protease activity sensors for use in live cells as well as in vitro assays and have validated their utility for antiviral drug discovery using GC376 as a proof of principle. The use of BRET, with NLuc as the bioluminescence donor and mNG as the resonance energy acceptor, enabled highly sensitive detection of M^pro^ proteolytic activity in live cells.Additionally, the sensors developed here did not show any cleavage, either in the absence of M^pro^ or in the presence of the catalytically dead, C145A mutant M^pro^, thus, displaying high specificity. The BRET-based sensors developed here appears to possess better sensitivity and specificity compared to the FlipGFP-based M^pro^ with respect to. The BRET-based sensor allowed detection of molecular crowding-mediated increased activity of M^pro^ enabling the discovery of reduced inhibitory efficacy of GC376 under a crowded condition. We believe that these sensors will find utility in both detecting active SARS-CoV-2 infection as well in screening antivirals developed for targeting M^pro^ proteolytic cleavage activity in live cells. Additionally,they can be utilized for determining effects of genetic variation in the M^pro^ amino acid sequence that may arise during the evolution of the virus.

## Supporting information

Short-sensor-linker-MD-traj-1000ns

Long-sensor-linker-MD-traj-1000ns

## Acknowledgements

This work is supported by an internal funding from the College of Health & Life Sciences, Hamad Bin Khalifa University, a member of the Qatar Foundation. A.M.G. is supported by a postdoctoral fellowship and W.S.A., S.R. A.G.N.F and S.M.N.U are supported by scholarships from the College of Health & Life Sciences, Hamad Bin Khalifa University, a member of the Qatar Foundation. Some of the computational research work reported in the manuscript were performed using high-performance computer resources and services provided by the Research Computing group in Texas A&M University at Qatar. Research Computing is funded by the Qatar Foundation for Education, Science and Community Development (http://www.qf.org.qa).

## Author Contributions

K.H.B. conceived the experiments. A.M.G, W.S.A, S.R, A.G.N.F, S.M.N.U, M.A. and

K.H.B. performed experiments, analyzed data, prepared figures, and wrote the manuscript. All authors reviewed the manuscript.

## Competing interests

K.H.B. & A.M.G. are inventors on a US Provisional Patent application number 63/275217, assigned to Qatar Foundation for Education, Science and Community Development, entitled “BRET-based coronavirus M^pro^ protease sensor and uses thereof” relating to the development and uses of the M^pro^ sensor. Other authors declare no competing interests.

## References

1. Lu, R.; Zhao, X.; Li, J.; Niu, P.; Yang, B.; Wu, H.; Wang, W.; Song, H.; Huang, B.; Zhu, N.; Bi, Y.; Ma, X.; Zhan, F.; Wang, L.; Hu, T.; Zhou, H.; Hu, Z.; Zhou, W.; Zhao, L.; Chen, J.; Meng, Y.; Wang, J.; Lin, Y.; Yuan, J.; Xie, Z.; Ma, J.; Liu, W. J.; Wang, D.; Xu, W.; Holmes, E. C.; Gao, G. F.; Wu, G.; Chen, W.; Shi, W.; Tan, W., Genomic characterisation and epidemiology of 2019 novel coronavirus: implications for virus origins and receptor binding. Lancet 2020, 395 (10224), 565–574.

2. Coelho, C.; Gallo, G.; Campos, C. B.; Hardy, L.; Wurtele, M., Biochemical screening for SARS-CoV-2 main protease inhibitors. PloS one 2020, 15 (10), e0240079.

3. Kelly, J. A.; Olson, A. N.; Neupane, K.; Munshi, S.; San Emeterio, J.; Pollack, L.; Woodside, M. T.; Dinman, J. D., Structural and functional conservation of the programmed -1 ribosomal frameshift signal of SARS coronavirus 2 (SARS-CoV-2). J. Biol. Chem. 2020, 295 (31), 10741–10748.

4. Jin, Z.; Du, X.; Xu, Y.; Deng, Y.; Liu, M.; Zhao, Y.; Zhang, B.; Li, X.; Zhang, L.; Peng, C.; Duan, Y.; Yu, J.; Wang, L.; Yang, K.; Liu, F.; Jiang, R.; Yang, X.; You, T.; Liu, X.; Yang, X.; Bai, F.; Liu, H.; Liu, X.; Guddat, L. W.; Xu, W.; Xiao, G.; Qin, C.; Shi, Z.; Jiang, H.; Rao, Z.; Yang, H., Structure of M(pro) from SARS-CoV-2 and discovery of its inhibitors. Nature 2020, 582 (7811), 289–293.

5. Zhang, L.; Lin, D.; Sun, X.; Curth, U.; Drosten, C.; Sauerhering, L.; Becker, S.; Rox, K.; Hilgenfeld, R., Crystal structure of SARS-CoV-2 main protease provides a basis for design of improved alpha-ketoamide inhibitors. Science 2020, 368 (6489), 409–412.

6. Naqvi, A. A. T.; Fatima, K.; Mohammad, T.; Fatima, U.; Singh, I. K.; Singh, A.; Atif, S. M.; Hariprasad, G.; Hasan, G. M.; Hassan, M. I., Insights into SARS-CoV-2 genome, structure, evolution, pathogenesis and therapies: Structural genomics approach. Biochim Biophys Acta Mol Basis Dis 2020, 1866 (10), 165878.

7. Cui, W.; Yang, K.; Yang, H., Recent Progress in the Drug Development Targeting SARS-CoV-2 Main Protease as Treatment for COVID-19. Frontiers in Molecular Biosciences 2020, 7 (398).

8. Yang, H.; Yang, M.; Ding, Y.; Liu, Y.; Lou, Z.; Zhou, Z.; Sun, L.; Mo, L.; Ye, S.; Pang, H.; Gao, G. F.; Anand, K.; Bartlam, M.; Hilgenfeld, R.; Rao, Z., The crystal structures of severe acute respiratory syndrome virus main protease and its complex with an inhibitor. Proc. Natl. Acad. Sci. U. S. A. 2003, 100 (23), 13190–13195.

9. Ramos-Guzmán, C. A.; Ruiz-Pernía, J. J.; Tuñón, I., Unraveling the SARS-CoV-2 Main Protease Mechanism Using Multiscale Methods. ACS Catalysis 2020, 10 (21), 12544–12554.

10. Rut, W.; Groborz, K.; Zhang, L.; Sun, X.; Zmudzinski, M.; Pawlik, B.; Wang, X.; Jochmans, D.; Neyts, J.; Mlynarski, W.; Hilgenfeld, R.; Drag, M., SARS-CoV-2 M(pro) inhibitors and activity-based probes for patient-sample imaging. Nat. Chem. Biol. 2020.

11. Xue, X.; Yang, H.; Shen, W.; Zhao, Q.; Li, J.; Yang, K.; Chen, C.; Jin, Y.; Bartlam, M.; Rao, Z., Production of authentic SARS-CoV M(pro) with enhanced activity: application as a novel tag-cleavage endopeptidase for protein overproduction. J. Mol. Biol. 2007, 366 (3), 965–75.

12. Du, Q. S.; Wang, S. Q.; Zhu, Y.; Wei, D. Q.; Guo, H.; Sirois, S.; Chou, K. C., Polyprotein cleavage mechanism of SARS CoV Mpro and chemical modification of the octapeptide. Peptides 2004, 25 (11), 1857–64.

13. Goetz, D. H.; Choe, Y.; Hansell, E.; Chen, Y. T.; McDowell, M.; Jonsson, C. B.; Roush, W. R.; McKerrow, J.; Craik, C. S., Substrate specificity profiling and identification of a new class of inhibitor for the major protease of the SARS coronavirus. Biochemistry 2007, 46 (30), 8744–52.

14. Zhang, L.; Lin, D.; Sun, X.; Curth, U.; Drosten, C.; Sauerhering, L.; Becker, S.; Rox, K.; Hilgenfeld, R., Crystal structure of SARS-CoV-2 main protease provides a basis for design of improved α-ketoamide inhibitors. Science 2020, eabb3405.

15. Roda, A.; Pasini, P.; Mirasoli, M.; Michelini, E.; Guardigli, M., Biotechnological applications of bioluminescence and chemiluminescence. Trends Biotechnol. 2004, 22 (6), 295–303.

16. New, D. C.; Miller-Martini, D. M.; Wong, Y. H., Reporter gene assays and their applications to bioassays of natural products. Phytother. Res. 2003, 17 (5), 439–48.

17. Bacart, J.; Corbel, C.; Jockers, R.; Bach, S.; Couturier, C., The BRET technology and its application to screening assays. Biotechnol J 2008, 3 (3), 311–24.

18. Biswas, K. H.; Sopory, S.; Visweswariah, S. S., The GAF domain of the cGMP-binding, cGMP-specific phosphodiesterase (PDE5) is a sensor and a sink for cGMP. Biochemistry 2008, 47 (11), 3534–43.

19. Elshabrawy, H. A.; Fan, J.; Haddad, C. S.; Ratia, K.; Broder, C. C.; Caffrey, M.; Prabhakar, B. S., Identification of a broad-spectrum antiviral small molecule against severe acute respiratory syndrome coronavirus and Ebola, Hendra, and Nipah viruses by using a novel high-throughput screening assay. J. Virol. 2014, 88 (8), 4353–65.

20. Kilianski, A.; Mielech, A. M.; Deng, X.; Baker, S. C., Assessing activity and inhibition of Middle East respiratory syndrome coronavirus papain-like and 3C-like proteases using luciferase-based biosensors. J. Virol. 2013, 87 (21), 11955–11962.

21. Zhou, J.; Fang, L.; Yang, Z.; Xu, S.; Lv, M.; Sun, Z.; Chen, J.; Wang, D.; Gao, J.; Xiao, S., Identification of novel proteolytically inactive mutations in coronavirus 3C-like protease using a combined approach. FASEB J. 2019, 33 (12), 14575–14587.

22. Zhu, W.; Xu, M.; Chen, C. Z.; Guo, H.; Shen, M.; Hu, X.; Shinn, P.; Klumpp-Thomas, C.; Michael, S. G.; Zheng, W., Identification of SARS-CoV-2 3CL Protease Inhibitors by a Quantitative High-Throughput Screening. ACS Pharmacol Transl Sci 2020, 3 (5), 1008–1016.

23. Tripathi, P. K.; Upadhyay, S.; Singh, M.; Raghavendhar, S.; Bhardwaj, M.; Sharma, P.; Patel, A. K., Screening and evaluation of approved drugs as inhibitors of main protease of SARS-CoV-2. Int. J. Biol. Macromol. 2020, 164, 2622–2631.

24. Brown, A. S.; Ackerley, D. F.; Calcott, M. J., High-Throughput Screening for Inhibitors of the SARS-CoV-2 Protease Using a FRET-Biosensor. Molecules 2020, 25 (20).

25. Ma, C.; Hu, Y.; Townsend, J. A.; Lagarias, P. I.; Marty, M. T.; Kolocouris, A.; Wang, J., Ebselen, Disulfiram, Carmofur, PX-12, Tideglusib, and Shikonin Are Nonspecific Promiscuous SARS-CoV-2 Main Protease Inhibitors. ACS Pharmacol Transl Sci 2020, 3 (6), 1265–1277.

26. Hung, H. C.; Ke, Y. Y.; Huang, S. Y.; Huang, P. N.; Kung, Y. A.; Chang, T. Y.; Yen, K. J.; Peng, T. T.; Chang, S. E.; Huang, C. T.; Tsai, Y. R.; Wu, S. H.; Lee, S. J.; Lin, J. H.; Liu, B. S.; Sung, W. C.; Shih, S. R.; Chen, C. T.; Hsu, J. T., Discovery of M Protease Inhibitors Encoded by SARS-CoV-2. Antimicrob Agents Chemother 2020, 64 (9).

27. Ma, C.; Sacco, M. D.; Hurst, B.; Townsend, J. A.; Hu, Y.; Szeto, T.; Zhang, X.; Tarbet, B.; Marty, M. T.; Chen, Y.; Wang, J., Boceprevir, GC-376, and calpain inhibitors II, XII inhibit SARS-CoV-2 viral replication by targeting the viral main protease. Cell Res 2020, 30 (8), 678–692.

28. Froggatt, H. M.; Heaton, B. E.; Heaton, N. S., Development of a Fluorescence-Based, High-Throughput SARS-CoV-2 3CL(pro) Reporter Assay. J. Virol. 2020, 94 (22).

29. Li, X.; Lidsky, P.; Xiao, Y.; Wu, C. T.; Garcia-Knight, M.; Yang, J.; Nakayama, T.; Nayak, J. V.; Jackson, P. K.; Andino, R.; Shu, X., Ethacridine inhibits SARS-CoV-2 by inactivating viral particles in cellular models. bioRxiv 2020.

30. Drayman, N.; Jones, K. A.; Azizi, S.-A.; Froggatt, H. M.; Tan, K.; Maltseva, N. I.; Chen, S.; Nicolaescu, V.; Dvorkin, S.; Furlong, K.; Kathayat, R. S.; Firpo, M. R.; Mastrodomenico, V.; Bruce, E. A.; Schmidt, M. M.; Jedrzejczak, R.; Muñoz-Alía, M. Á.; Schuster, B.; Nair, V.; Botten, J. W.; Brooke, C. B.; Baker, S. C.; Mounce, B. C.; Heaton, N. S.; Dickinson, B. C.; Jaochimiak, A.; Randall, G.; Tay, S., Drug repurposing screen identifies masitinib as a 3CLpro inhibitor that blocks replication of SARS-CoV-2 in vitro. bioRxiv : the preprint server for biology 2020, 2020.08.31.274639.

31. Heroux, M.; Breton, B.; Hogue, M.; Bouvier, M., Assembly and signaling of CRLR and RAMP1 complexes assessed by BRET. Biochemistry 2007, 46 (23), 7022–33.

32. Nikolaev, V. O.; Gambaryan, S.; Lohse, M. J., Fluorescent sensors for rapid monitoring of intracellular cGMP. Nat Methods 2006, 3 (1), 23–5.

33. Min, S.-H. F. A.R.; Trull, K.J.; Tat, K.; Varney, S.A.; Tantama, M., Ratiometric BRET Measurements of ATP with a Genetically-Encoded Luminescent Sensor. Sensors 2019, 19 (16), 3502.

34. Rathod, M.; Mal, A.; De, A., Reporter-Based BRET Sensors for Measuring Biological Functions In Vivo. Methods Mol. Biol. 2018, 1790, 51–74.

35. Xie, Q.; Soutto, M.; Xu, X.; Zhang, Y.; Johnson, C. H., Bioluminescence resonance energy transfer (BRET) imaging in plant seedlings and mammalian cells. Methods Mol. Biol. 2011, 680, 3–28.

36. Biswas, K. H.; Visweswariah, S. S., Distinct allostery induced in the cyclic GMP-binding, cyclic GMP-specific phosphodiesterase (PDE5) by cyclic GMP, sildenafil, and metal ions. J Biol Chem 2011, 286 (10), 8545–54.

37. Biswas, K.; Sopory, S.; Visweswariah, S., The GAF domain of the cGMP-binding, cGMP-specific phosphodiesterase (PDE5) is a sensor and a sink for cGMP. Biochemistry 2008, 47 (11), 3534.

38. Biswas, K. H.; Badireddy, S.; Rajendran, A.; Anand, G. S.; Visweswariah, S. S., Cyclic nucleotide binding and structural changes in the isolated GAF domain of Anabaena adenylyl cyclase, CyaB2. PeerJ 2015, 3, e882.

39. Biswas, K. H.; Visweswariah, S. S., Buffer NaCl concentration regulates Renilla luciferase activity and ligand-induced conformational changes in the BRET-based PDE5 sensor. Matters 2017, 10.19185/matters.201702000015.

40. Chien, Y. H.; Jiang, N.; Li, F.; Zhang, F.; Zhu, C.; Leckband, D., Two stage cadherin kinetics require multiple extracellular domains but not the cytoplasmic region. J. Biol. Chem. 2008, 283 (4), 1848–56.

41. Biswas, K. H.; Visweswariah, S. S., Distinct allostery induced in the cyclic GMP-binding, cyclic GMP-specific phosphodiesterase (PDE5) by cyclic GMP, sildenafil, and metal ions. J. Biol. Chem. 2011, 286 (10), 8545–54.

42. Heffern, M. C., Diversifying the Glowing Bioluminescent Toolbox. ACS Cent Sci 2017, 3 (12), 1234–1236.

43. Clavel, D.; Gotthard, G.; von Stetten, D.; De Sanctis, D.; Pasquier, H.; Lambert, G. G.; Shaner, N. C.; Royant, A., Structural analysis of the bright monomeric yellow-green fluorescent protein mNeonGreen obtained by directed evolution. Acta Crystallogr D Struct Biol 2016, 72 (Pt 12), 1298–1307.

44. Hostettler, L.; Grundy, L.; Kaser-Pebernard, S.; Wicky, C.; Schafer, W. R.; Glauser, D. A., The Bright Fluorescent Protein mNeonGreen Facilitates Protein Expression Analysis In Vivo. G3 (Bethesda) 2017, 7 (2), 607–615.

45. Steiert, F.; Petrov, E. P.; Schultz, P.; Schwille, P.; Weidemann, T., Photophysical Behavior of mNeonGreen, an Evolutionarily Distant Green Fluorescent Protein. Biophys. J. 2018, 114 (10), 2419–2431.

46. Tanida-Miyake, E.; Koike, M.; Uchiyama, Y.; Tanida, I., Optimization of mNeonGreen for Homo sapiens increases its fluorescent intensity in mammalian cells. PloS one 2018, 13 (1), e0191108.

47. Dale, N. C.; Johnstone, E. K. M.; White, C. W.; Pfleger, K. D. G., NanoBRET: The Bright Future of Proximity-Based Assays. Front Bioeng Biotechnol 2019, 7, 56.

48. Hall, M. P.; Unch, J.; Binkowski, B. F.; Valley, M. P.; Butler, B. L.; Wood, M. G.; Otto, P.; Zimmerman, K.; Vidugiris, G.; Machleidt, T.; Robers, M. B.; Benink, H. A.; Eggers, C. T.; Slater, M. R.; Meisenheimer, P. L.; Klaubert, D. H.; Fan, F.; Encell, L. P.; Wood, K. V., Engineered luciferase reporter from a deep sea shrimp utilizing a novel imidazopyrazinone substrate. ACS Chem Biol 2012, 7 (11), 1848–57.

49. den Hamer, A.; Dierickx, P.; Arts, R.; de Vries, J.; Brunsveld, L.; Merkx, M., Bright Bioluminescent BRET Sensor Proteins for Measuring Intracellular Caspase Activity. ACS Sens 2017, 2 (6), 729–734.

50. Zarowny, L.; Aggarwal, A.; Rutten, V. M. S.; Kolb, I.; Project, G.; Patel, R.; Huang, H. Y.; Chang, Y. F.; Phan, T.; Kanyo, R.; Ahrens, M. B.; Allison, W. T.; Podgorski, K.; Campbell, R. E., Bright and High-Performance Genetically Encoded Ca(2+) Indicator Based on mNeonGreen Fluorescent Protein. ACS Sens 2020, 5 (7), 1959–1968.

51. Aird, E. J.; Tompkins, K. J.; Ramirez, M. P.; Gordon, W. R., Enhanced Molecular Tension Sensor Based on Bioluminescence Resonance Energy Transfer (BRET). ACS Sens 2020, 5 (1), 34–39.

52. Oishi, A.; Dam, J.; Jockers, R., beta-Arrestin-2 BRET Biosensors Detect Different beta-Arrestin-2 Conformations in Interaction with GPCRs. ACS Sens 2020, 5 (1), 57–64.

53. Parag-Sharma, K.; O’Banion, C. P.; Henry, E. C.; Musicant, A. M.; Cleveland, J. L.; Lawrence, D. S.; Amelio, A. L., Engineered BRET-Based Biologic Light Sources Enable Spatiotemporal Control over Diverse Optogenetic Systems. ACS Synth Biol 2020, 9 (1), 1–9.

54. Machleidt, T.; Woodroofe, C. C.; Schwinn, M. K.; Mendez, J.; Robers, M. B.; Zimmerman, K.; Otto, P.; Daniels, D. L.; Kirkland, T. A.; Wood, K. V., NanoBRET--A Novel BRET Platform for the Analysis of Protein-Protein Interactions. ACS Chem Biol 2015, 10 (8), 1797–804.

55. Stoddart, L. A.; Kilpatrick, L. E.; Hill, S. J., NanoBRET Approaches to Study Ligand Binding to GPCRs and RTKs. Trends Pharmacol. Sci. 2018, 39 (2), 136–147.

56. Weihs, F.; Wang, J.; Pfleger, K. D. G.; Dacres, H., Experimental determination of the bioluminescence resonance energy transfer (BRET) Forster distances of NanoBRET and red-shifted BRET pairs. Anal Chim Acta X 2020, 6, 100059.

57. Gordon, D. E.; Jang, G. M.; Bouhaddou, M.; Xu, J.; Obernier, K.; White, K. M.; O’Meara, M. J.; Rezelj, V. V.; Guo, J. Z.; Swaney, D. L.; Tummino, T. A.; Huettenhain, R.; Kaake, R. M.; Richards, A. L.; Tutuncuoglu, B.; Foussard, H.; Batra, J.; Haas, K.; Modak, M.; Kim, M.; Haas, P.; Polacco, B. J.; Braberg, H.; Fabius, J. M.; Eckhardt, M.; Soucheray, M.; Bennett, M. J.; Cakir, M.; McGregor, M. J.; Li, Q.; Meyer, B.; Roesch, F.; Vallet, T.; Mac Kain, A.; Miorin, L.; Moreno, E.; Naing, Z. Z. C.; Zhou, Y.; Peng, S.; Shi, Y.; Zhang, Z.; Shen, W.; Kirby, I. T.; Melnyk, J. E.; Chorba, J. S.; Lou, K.; Dai, S. A.; Barrio-Hernandez, I.; Memon, D.; Hernandez-Armenta, C.; Lyu, J.; Mathy, C. J. P.; Perica, T.; Pilla, K. B.; Ganesan, S. J.; Saltzberg, D. J.; Rakesh, R.; Liu, X.; Rosenthal, S. B.; Calviello, L.; Venkataramanan, S.; Liboy-Lugo, J.; Lin, Y.; Huang, X. P.; Liu, Y.; Wankowicz, S. A.; Bohn, M.; Safari, M.; Ugur, F. S.; Koh, C.; Savar, N. S.; Tran, Q. D.; Shengjuler, D.; Fletcher, S. J.; O’Neal, M. C.; Cai, Y.; Chang, J. C. J.; Broadhurst, D. J.; Klippsten, S.; Sharp, P. P.; Wenzell, N. A.; Kuzuoglu, D.; Wang, H. Y.; Trenker, R.; Young, J. M.; Cavero, D. A.; Hiatt, J.; Roth, T. L.; Rathore, U.; Subramanian, A.; Noack, J.; Hubert, M.; Stroud, R. M.; Frankel, A. D.; Rosenberg, O. S.; Verba, K. A.; Agard, D. A.; Ott, M.; Emerman, M.; Jura, N.; von Zastrow, M.; Verdin, E.; Ashworth, A.; Schwartz, O.; d’Enfert, C.; Mukherjee, S.; Jacobson, M.; Malik, H. S.; Fujimori, D. G.; Ideker, T.; Craik, C. S.; Floor, S. N.; Fraser, J. S.; Gross, J. D.; Sali, A.; Roth, B. L.; Ruggero, D.; Taunton, J.; Kortemme, T.; Beltrao, P.; Vignuzzi, M.; Garcia-Sastre, A.; Shokat, K. M.; Shoichet, B. K.; Krogan, N. J., A SARS-CoV-2 protein interaction map reveals targets for drug repurposing. Nature 2020.

58. Huang, C.; Wei, P.; Fan, K.; Liu, Y.; Lai, L., 3C-like proteinase from SARS coronavirus catalyzes substrate hydrolysis by a general base mechanism. Biochemistry 2004, 43 (15), 4568–74.

59. Shan, Y. F.; Li, S. F.; Xu, G. J., A novel auto-cleavage assay for studying mutational effects on the active site of severe acute respiratory syndrome coronavirus 3C-like protease. Biochem Biophys Res Commun 2004, 324 (2), 579–83.

60. Fu, L.; Ye, F.; Feng, Y.; Yu, F.; Wang, Q.; Wu, Y.; Zhao, C.; Sun, H.; Huang, B.; Niu, P.; Song, H.; Shi, Y.; Li, X.; Tan, W.; Qi, J.; Gao, G. F., Both Boceprevir and GC376 efficaciously inhibit SARS-CoV-2 by targeting its main protease. Nat. Commun. 2020, 11 (1), 4417.

61. Dai, W.; Zhang, B.; Jiang, X. M.; Su, H.; Li, J.; Zhao, Y.; Xie, X.; Jin, Z.; Peng, J.; Liu, F.; Li, C.; Li, Y.; Bai, F.; Wang, H.; Cheng, X.; Cen, X.; Hu, S.; Yang, X.; Wang, J.; Liu, X.; Xiao, G.; Jiang, H.; Rao, Z.; Zhang, L. K.; Xu, Y.; Yang, H.; Liu, H., Structure-based design of antiviral drug candidates targeting the SARS-CoV-2 main protease. Science 2020, 368 (6497), 1331–1335.

62. Sacco, M. D.; Ma, C.; Lagarias, P.; Gao, A.; Townsend, J. A.; Meng, X.; Dube, P.; Zhang, X.; Hu, Y.; Kitamura, N.; Hurst, B.; Tarbet, B.; Marty, M. T.; Kolocouris, A.; Xiang, Y.; Chen, Y.; Wang, J., Structure and inhibition of the SARS-CoV-2 main protease reveal strategy for developing dual inhibitors against M<sup>pro</sup> and cathepsin L. Science Advances 2020, 6 (50), eabe0751.

63. Katoh, K.; Rozewicki, J.; Yamada, K. D., MAFFT online service: multiple sequence alignment, interactive sequence choice and visualization. Brief. Bioinform. 2019, 20 (4), 1160–1166.

64. Kuraku, S.; Zmasek, C. M.; Nishimura, O.; Katoh, K., aLeaves facilitates on-demand exploration of metazoan gene family trees on MAFFT sequence alignment server with enhanced interactivity. Nucleic Acids Res. 2013, 41 (Web Server issue), W22–8.

65. Xue, X.; Yu, H.; Yang, H.; Xue, F.; Wu, Z.; Shen, W.; Li, J.; Zhou, Z.; Ding, Y.; Zhao, Q.; Zhang, X. C.; Liao, M.; Bartlam, M.; Rao, Z., Structures of two coronavirus main proteases: implications for substrate binding and antiviral drug design. J. Virol. 2008, 82 (5), 2515–27.

66. Eswar, N.; Eramian, D.; Webb, B.; Shen, M.-Y.; Sali, A., Protein Structure Modeling With MODELLER. Methods in molecular biology (Clifton, NJ) 2008, 426, 145–159.

67. Laskowski, R. A.; MacArthur, M. W.; Moss, D. S.; Thornton, J. M., PROCHECK: a program to check the stereochemical quality of protein structures. Journal of applied crystallography 1993, 26 (2), 283–291.

68. Jo, S.; Kim, T.; Iyer, V. G.; Im, W., CHARMM-GUI: a web-based graphical user interface for CHARMM. Journal of computational chemistry 2008, 29 (11), 1859–1865.

69. Jorgensen, W. L.; Chandrasekhar, J.; Madura, J. D.; Impey, R. W.; Klein, M. L., Comparison of simple potential functions for simulating liquid water. The Journal of chemical physics 1983, 79 (2), 926–935.

70. Phillips, J. C.; Braun, R.; Wang, W.; Gumbart, J.; Tajkhorshid, E.; Villa, E.; Chipot, C.; Skeel, R. D.; Kalé, L.; Schulten, K., Scalable Molecular Dynamics With NAMD. Journal of computational chemistry 2005, 26 (16), 1781–1802.

71. Huang, J.; Rauscher, S.; Nawrocki, G.; Ran, T.; Feig, M.; de Groot, B. L.; Grubmüller, H.; MacKerell Jr, A. D., CHARMM36m: an improved force field for folded and intrinsically disordered proteins. Nature methods 2017, 14 (1), 71–73.

72. Feller, S. E.; Zhang, Y.; Pastor, R. W.; Brooks, B. R., Constant pressure molecular dynamics simulation: the Langevin piston method. The Journal of chemical physics 1995, 103 (11), 4613–4621.

73. Pang, Y. T.; Miao, Y.; Wang, Y.; McCammon, J. A., Gaussian Accelerated Molecular Dynamics in NAMD. Journal of Chemical Theory and Computation 2017, 13 (1), 9.

74. Miao, Y.; Feixas, F.; Eun, C.; McCammon, J. A., Accelerated Molecular Dynamics Simulations of Protein Folding. Journal of computational chemistry 2015, 36 (20), 1536.

75. Zhang, Q.; Tan, S.; Xiao, T.; Liu, H.; Shah, S. J. A.; Liu, H., Probing the Molecular Mechanism of Rifampin Resistance Caused by the Point Mutations S456L and D441V on Mycobacterium Tuberculosis RNA Polymerase Through Gaussian Accelerated Molecular Dynamics Simulation. Antimicrobial agents and chemotherapy 2020, 64 (7), e02476–19.

76. Miao, Y.; Feher, V. A.; McCammon, J. A., Gaussian Accelerated Molecular Dynamics: Unconstrained Enhanced Sampling and Free Energy Calculation. Journal of Chemical Theory and Computation 2015, 11 (8), 3584.

77. Miao, Y.; McCammon, J. A., Gaussian Accelerated Molecular Dynamics: Theory, Implementation, and Applications. Annual reports in computational chemistry 2017, 13, 231.

78. Humphrey, W.; Dalke, A.; Schulten, K., VMD: visual molecular dynamics. Journal of molecular graphics 1996, 14 (1), 33.

79. Saifaldeen, M.; Al-Ansari, D. E.; Ramotar, D.; Aouida, M., Dead Cas9-sgRNA Complex Shelters Vulnerable DNA Restriction Enzyme Sites from Cleavage for Cloning Applications. CRISPR J 2021, 4 (2), 275–289.

80. Saha, S.; Biswas, K. H.; Kondapalli, C.; Isloor, N.; Visweswariah, S. S., The linker region in receptor guanylyl cyclases is a key regulatory module: mutational analysis of guanylyl cyclase C. J. Biol. Chem. 2009, 284 (40), 27135–45.

81. Fiskerstrand, T.; Arshad, N.; Haukanes, B. I.; Tronstad, R. R.; Pham, K. D.; Johansson, S.; Havik, B.; Tonder, S. L.; Levy, S. E.; Brackman, D.; Boman, H.; Biswas, K. H.; Apold, J.; Hovdenak, N.; Visweswariah, S. S.; Knappskog, P. M., Familial diarrhea syndrome caused by an activating GUCY2C mutation. N. Engl. J. Med. 2012, 366 (17), 1586–95.

82. Biswas, K. H.; Hartman, K. L.; Yu, C. H.; Harrison, O. J.; Song, H.; Smith, A. W.; Huang, W. Y.; Lin, W. C.; Guo, Z.; Padmanabhan, A.; Troyanovsky, S. M.; Dustin, M. L.; Shapiro, L.; Honig, B.; Zaidel-Bar, R.; Groves, J. T., E-cadherin junction formation involves an active kinetic nucleation process. Proc Natl Acad Sci U S A 2015, 112 (35), 10932–7.

83. Biswas, K. H.; Hartman, K. L.; Zaidel-Bar, R.; Groves, J. T., Sustained alpha-catenin Activation at E-cadherin Junctions in the Absence of Mechanical Force. Biophys. J. 2016, 111 (5), 1044–52.

84. Biswas, K. H.; Zhongwen, C.; Dubey, A. K.; Oh, D.; Groves, J. T., Multicomponent Supported Membrane Microarray for Monitoring Spatially Resolved Cellular Signaling Reactions. Adv. Biosyst. 2018, 2 (4), 1800015.

85. Gordon, D. E.; Jang, G. M.; Bouhaddou, M.; Xu, J.; Obernier, K.; O’Meara, M. J.; Guo, J. Z.; Swaney, D. L.; Tummino, T. A.; Huttenhain, R.; Kaake, R. M.; Richards, A. L.; Tutuncuoglu, B.; Foussard, H.; Batra, J.; Haas, K.; Modak, M.; Kim, M.; Haas, P.; Polacco, B. J.; Braberg, H.; Fabius, J. M.; Eckhardt, M.; Soucheray, M.; Bennett, M. J.; Cakir, M.; McGregor, M. J.; Li, Q.; Naing, Z. Z. C.; Zhou, Y.; Peng, S.; Kirby, I. T.; Melnyk, J. E.; Chorba, J. S.; Lou, K.; Dai, S. A.; Shen, W.; Shi, Y.; Zhang, Z.; Barrio-Hernandez, I.; Memon, D.; Hernandez-Armenta, C.; Mathy, C. J. P.; Perica, T.; Pilla, K. B.; Ganesan, S. J.; Saltzberg, D. J.; Ramachandran, R.; Liu, X.; Rosenthal, S. B.; Calviello, L.; Venkataramanan, S.; Lin, Y.; Wankowicz, S. A.; Bohn, M.; Trenker, R.; Young, J. M.; Cavero, D.; Hiatt, J.; Roth, T.; Rathore, U.; Subramanian, A.; Noack, J.; Hubert, M.; Roesch, F.; Vallet, T.; Meyer, B.; White, K. M.; Miorin, L.; Agard, D.; Emerman, M.; Ruggero, D.; Garcia-Sastre, A.; Jura, N.; von Zastrow, M.; Taunton, J.; Schwartz, O.; Vignuzzi, M.; d’Enfert, C.; Mukherjee, S.; Jacobson, M.; Malik, H. S.; Fujimori, D. G.; Ideker, T.; Craik, C. S.; Floor, S.; Fraser, J. S.; Gross, J.; Sali, A.; Kortemme, T.; Beltrao, P.; Shokat, K.; Shoichet, B. K.; Krogan, N. J., A SARS-CoV-2-Human Protein-Protein Interaction Map Reveals Drug Targets and Potential Drug-Repurposing. bioRxiv 2020.

86. Schindelin, J.; Arganda-Carreras, I.; Frise, E.; Kaynig, V.; Longair, M.; Pietzsch, T.; Preibisch, S.; Rueden, C.; Saalfeld, S.; Schmid, B.; Tinevez, J. Y.; White, D. J.; Hartenstein, V.; Eliceiri, K.; Tomancak, P.; Cardona, A., Fiji: an open-source platform for biological-image analysis. Nat Methods 2012, 9 (7), 676–82.

87. Grum-Tokars, V.; Ratia, K.; Begaye, A.; Baker, S. C.; Mesecar, A. D., Evaluating the 3C-like protease activity of SARS-Coronavirus: recommendations for standardized assays for drug discovery. Virus Res 2008, 133 (1), 63–73.

88. Bartke, T.; Pohl, C.; Pyrowolakis, G.; Jentsch, S., Dual role of BRUCE as an antiapoptotic IAP and a chimeric E2/E3 ubiquitin ligase. Mol Cell 2004, 14 (6), 801–11.

89. Dacres, H.; Dumancic, M. M.; Horne, I.; Trowell, S. C., Direct comparison of fluorescence-and bioluminescence-based resonance energy transfer methods for real-time monitoring of thrombin-catalysed proteolytic cleavage. Biosens Bioelectron 2009, 24 (5), 1164–70.

90. Chuck, C. P.; Chow, H. F.; Wan, D. C.; Wong, K. B., Profiling of substrate specificities of 3C-like proteases from group 1, 2a, 2b, and 3 coronaviruses. PloS one 2011, 6 (11), e27228.

91. Fan, K.; Ma, L.; Han, X.; Liang, H.; Wei, P.; Liu, Y.; Lai, L., The substrate specificity of SARS coronavirus 3C-like proteinase. Biochem Biophys Res Commun 2005, 329 (3), 934–40.

92. Chuck, C. P.; Chong, L. T.; Chen, C.; Chow, H. F.; Wan, D. C.; Wong, K. B., Profiling of substrate specificity of SARS-CoV 3CL. PloS one 2010, 5 (10), e13197.

93. Phillips, J. C.; Braun, R.; Wang, W.; Gumbart, J.; Tajkhorshid, E.; Villa, E.; Chipot, C.; Skeel, R. D.; Kalé, L.; Schulten, K., Scalable Molecular Dynamics with NAMD. J. Comput. Chem. 2005, 26 (16), 1781.

94. Balleza, E.; Kim, J. M.; Cluzel, P., Systematic characterization of maturation time of fluorescent proteins in living cells. Nat Methods 2018, 15 (1), 47–51.

95. Westerhausen, S.; Nowak, M.; Torres-Vargas, C. E.; Bilitewski, U.; Bohn, E.; Grin, I.; Wagner, S., A NanoLuc luciferase-based assay enabling the real-time analysis of protein secretion and injection by bacterial type III secretion systems. Mol. Microbiol. 2020, 113 (6), 1240–1254.

96. Kuo, C.-J.; Chi, Y.-H.; Hsu, J. T. A.; Liang, P.-H., Characterization of SARS main protease and inhibitor assay using a fluorogenic substrate. Biochemical and biophysical research communications 2004, 318 (4), 862–867.

97. Barrila, J.; Gabelli, S. B.; Bacha, U.; Amzel, L. M.; Freire, E., Mutation of Asn28 disrupts the dimerization and enzymatic activity of SARS 3CL(pro). Biochemistry 2010, 49 (20), 4308–17.

98. Tomar, S.; Johnston, M. L.; St John, S. E.; Osswald, H. L.; Nyalapatla, P. R.; Paul, L. N.; Ghosh, A. K.; Denison, M. R.; Mesecar, A. D., Ligand-induced Dimerization of Middle East Respiratory Syndrome (MERS) Coronavirus nsp5 Protease (3CLpro): IMPLICATIONS FOR nsp5 REGULATION AND THE DEVELOPMENT OF ANTIVIRALS. J. Biol. Chem. 2015, 290 (32), 19403–22.

99. Stagg, L.; Zhang, S. Q.; Cheung, M. S.; Wittung-Stafshede, P., Molecular crowding enhances native structure and stability of alpha/beta protein flavodoxin. Proc. Natl. Acad. Sci. U. S. A. 2007, 104 (48), 18976–81.

100. Uversky, V. N., Intrinsically disordered proteins in overcrowded milieu: Membrane-less organelles, phase separation, and intrinsic disorder. Current opinion in structural biology 2017, 44, 18–30.

101. Chebotareva, N. A.; Kurganov, B. I.; Livanova, N. B., Biochemical effects of molecular crowding. Biochemistry (Mosc.) 2004, 69 (11), 1239–51.

102. Maximova, K.; Wojtczak, J.; Trylska, J., Enzymatic activity of human immunodeficiency virus type 1 protease in crowded solutions. Eur. Biophys. J. 2019, 48 (7), 685–689.

103. Okamoto, D. N.; Oliveira, L. C.; Kondo, M. Y.; Cezari, M. H.; Szeltner, Z.; Juhasz, T.; Juliano, M. A.; Polgar, L.; Juliano, L.; Gouvea, I. E., Increase of SARS-CoV 3CL peptidase activity due to macromolecular crowding effects in the milieu composition. Biol. Chem. 2010, 391 (12), 1461–8.

104. Rivas, G.; Minton, A. P., Macromolecular Crowding In Vitro, In Vivo, and In Between. Trends in Biochemical Sciences 2016, 41 (11), 970–981.

105. Popielec, A.; Ostrowska, N.; Wojciechowska, M.; Feig, M.; Trylska, J., Crowded environment affects the activity and inhibition of the NS3/4A protease. Biochimie 2020, 176, 169–180.

106. Phillip, Y.; Sherman, E.; Haran, G.; Schreiber, G., Common crowding agents have only a small effect on protein-protein interactions. Biophys. J. 2009, 97 (3), 875–885.

107. Jia, M.; Sun, G.-Y.; Zhao, Y. X.; Liu, Z.-S.; Aisa, H. A., Effect of polyethylene glycol as a molecular crowding agent on reducing template consumption for preparation of molecularly imprinted polymers. Analytical Methods 2016, 8 (23), 4554–4562.

