## Supplementary material for "A Genetically encoded BRET-based SARS-CoV-2 Mpro protease activity sensor": Supp-info

### Supporting text & methods

**BRET-based $M^{pro}$ sensor construct details:**

*mNG-$M^{pro}$-Nter-auto-NLuc*:

Sequence:

MGSSHHHHHHSSGLVPRGSHMVSKGEEDNMASLPATHELHIFGSINGVDFDMVGQGTGNPNDGYEELNLKSTKGDLQ
FSPWILVPHIGYGFHQYLPYPDGMSPFQAAMVDGSGYQVHRTMQFEDGASLTVNYRYTYEGSHIKGEAQVKGTGFPA
DGPVMTNSLTAADWCRSKKTYPNDKTIISTFKWSYTTGNGKRYRSTARTTYTFAKPMAANYLKNQPMYVFRKTELKH
SKTELNFKEWQKAFTDVMGMDELYKEFGTENLYAVLQSGFRGSGGSMVFTLEDFVGDWRQTAGYNLDQVLEQGGVSS
LFQNLGVSVTPIQRIVLSGENGLKIDIHVIIPYEGLSGDQMGQIEKIFKVVYPVDDHHFKVILHYGTLVIDGVTPNM
IDYFGRPYEGIAVFDGKKITVTGTLWNGNKIIDERLINPDGSLLFRVTINGVTGWRLCERILA*

Predicted pI: 6.31
Predicted Mol. Wt.: 50048.54 (~50 kDa)
Predicted Mol. Wt. after cleavage (His-tagged fragment): Theoretical pI/Mw: 6.92 / 30178.78 (30 kDa)

*mNG-$M^{pro}$-Nter-auto-L-NLuc*:

Sequence:

MGSSHHHHHHSSGLVPRGSHMVSKGEEDNMASLPATHELHIFGSINGVDFDMVGQGTGNPNDGYEELNLKSTKGDLQ
FSPWILVPHIGYGFHQYLPYPDGMSPFQAAMVDGSGYQVHRTMQFEDGASLTVNYRYTYEGSHIKGEAQVKGTGFPA
DGPVMTNSLTAADWCRSKKTYPNDKTIISTFKWSYTTGNGKRYRSTARTTYTFAKPMAANYLKNQPMYVFRKTELKH
SKTELNFKEWQKAFTDVMGMDELYKEFGTENLYKTSAVLQSGFRKMEGSGGSMVFTLEDFVGDWRQTAGYNLDQVLE
QGGVSSLFQNLGVSVTPIQRIVLSGENGLKIDIHVIIPYEGLSGDQMGQIEKIFKVVYPVDDHHFKVILHYGTLVID
GVTPNMIDYFGRPYEGIAVFDGKKITVTGTLWNGNKIIDERLINPDGSLLFRVTINGVTGWRLCERILA*

Predicted pI: 6.41
Predicted Mol. Wt.: 50753.38 (~51 kDa)

Predicted Mol. Wt. after cleavage (His-tagged fragment): Theoretical pI/Mw: 7.26 / 30495.14

*Key*:

Yellow: His-tag
Green: mNeonGreen
Grey: Linkers
Red: cleavage sequence
Cyan: NLuc

**$M^{pro}$ protease constructs details:**

*WT $M^{pro}$ Sequence:*

As expressed from the pLVX-EF1alpha-SARS-CoV-2-nsp5-2xStrep-IRES-Puro[1] plasmid

MSGFRKMAFPSGKVEGCMVQVTCGTTTLNGLWLDDVVYCPRHVICTSEDMLNPNYEDLLIRKSN

HNFLVQAGNVQLRVIGHSMQNCVLKLKVDTANPKTPKYKFVRIQPGQTFSVLACYNGSPSGVYQ

CAMRPNFTIKGSFLNGSCGSVGFNIDYDCVSFCYMHHMELPTGVHAGTDLEGNFYGPFVDRQTA

QAAGTDTTITVNVLAWLYAAVINGDRWFLNRFTTTLNDFNLVAMKYNYEPLTQDHVDILGPLSA

QTGIAVLDMCASLKELLQNGMNGRTILGSALLEDEFTPFDVVRQCSGVTFQLEGGGGWSHPQFE

KGGGSGGGSGGGSWSHPQFEK

*C145A mutant $M^{pro}$ Sequence:*

As expressed from the pLVX-EF1alpha-SARS-CoV-2-nsp5-C145A-2xStrep-IRES-Puro[1] plasmid

MSGFRKMAFPSGKVEGCMVQVTCGTTTLNGLWLDDVVYCPRHVICTSEDMLNPNYEDLLIRKSN

HNFLVQAGNVQLRVIGHSMQNCVLKLKVDTANPKTPKYKFVRIQPGQTFSVLACYNGSPSGVYQ

CAMRPNFTIKGSFLNGSAGSVGFNIDYDCVSFCYMHHMELPTGVHAGTDLEGNFYGPFVDRQTA

QAAGTDTTITVNVLAWLYAAVINGDRWFLNRFTTTLNDFNLVAMKYNYEPLTQDHVDILGPLSA

QTGIAVLDMCASLKELLQNGMNGRTILGSALLEDEFTPFDVVRQCSGVTFQLEGGGGWSHPQFE
KGGGSGGGSGGGSWSHPQFEK

Key:
Green: M$^{pro}$
Yellow: Strep-tag II
Red: C145A mutation

**Dynamic Light Scattering (DLS) measurements**

DLS measurements of recombinantly purified proteins were performed using the Zetasizer Nano ZS (Malvern Panalytical, Malvern, United Kingdom). Proteins, either bovine serum albumin (BSA; Tocris Bioscience, Cat. No. 5217) or the short M$^{pro}$ sensor, were prepared in 1× TBS at a final concentration of 1 μM and 400 nM, respectively and light scattering measurements were performed. The purified short M$^{pro}$ biosensor protein was centrifuged prior to measurement at 14000 rpm for 1 hour at 4 °C and supernatant taken to remove any aggregates. Multiple size spectra ($N = 40$ for BSA and 61 for the short M$^{pro}$ biosensor) obtained from triplicate measurements for 5 s were used for the determination of average molecular size of the proteins.

**Script used for determining FlipGFP-based M$^{pro}$ sensor activity in live cells**

The following script was used for determining the percentage of GPF positive (GFP$^{+}$), number of transfected cells per image frame and total number of cells analyzed for each time points.

```
dir_img = getDirectory("Choose your directory of images");
list_img = getFileList(dir_img);

setBatchMode(true);

for (i=0; i<list_img.length; i++) {
```

```
open(dir_img + list_img[i]);

name=getTitle;
run("Split Channels");

selectWindow("C1-" +name);
run("Duplicate...", "title=duplicate");

selectWindow("duplicate");
run("Subtract Background...", "rolling=5");
run("Enhance Contrast...", "saturated=0.3");
run("Gaussian Blur...", "sigma=1");
run("Auto Local Threshold", "method=Bernsen radius=5 parameter_1=0
parameter_2=0 white");
run("Analyze Particles...", "size=5-Infinity circularity=0.10-1.00
add");

selectWindow("duplicate");
run("Close");

selectWindow("C1-" +name);
roiManager("Measure");
//print("C1-" +name);
selectWindow("C2-" +name);
roiManager("Measure");

     selectWindow("Results");
     saveAs("Text", dir_img+name+".txt");
     run("Close");

roiManager("Delete");

selectWindow("C1-" +name);
run("Close");

selectWindow("C2-" +name);
run("Close");
}
run("Close All");
```

**Bacterial expression and purification of $M^{pro}$-Nter-auto sensor**

The sensor was expressed in *Escherichia coli* (*E. coli*) BL21-CodonPlus cells (Agilent Technologies) in 100 mL of LB medium, as described previously[2]. Protein expression was induced by the addition of 0.5 mM isopropyl-β-D-thiogalactopyranoside (IPTG), followed by overnight incubation at 20℃. After harvesting the cells by centrifugation (10000 g, 10 min, 4℃), the pellet was resuspended in lysis buffer (10 mL per gram cell pellet; 10 mM phosphate buffer, 2.7 mM KCl, 507 mM NaCl, 10% glycerol, 20 mM imidazole and 0.1 mM DTT), followed by sonication. The supernatant was collected after centrifugation (18000 g, 90 min, 45℃). The sensor construct was purified using Ni-NTA affinity chromatography. The concentration of the sensor was determined using Bradford assay[3].

**In vitro enzyme kinetics measurements**

To determine the initial reaction velocity, a range of the short $M^{pro}$ sensor concentrations were incubated with 200 and 500 nM of the recombinantly purified $M^{pro}$ protein in a buffer containing 50 mM HEPES, 50 mM NaCl, 0.1% Triton X-100, 1 mM 1 mM Dithiothreitol (DTT) & 1 mM ethylenediamine tetraacetic acid (EDTA) in a total volume of 50 µL for 2.25 h at 37℃. Following incubation, each reaction was diluted to a final concentration of 20 nM of the $M^{pro}$ sensor and BRET measured using a Tecan SPARK® multimode microplate reader after addition of NLuc substrate. For initial reaction velocity calculation at each sensor (substrate) concentration, a background BRET value of 0.25 (obtained using NLuc alone) was subtracted from the initial as well as final BRET values. Reaction rates were then calculated as:

$$Rate = {[(BRET^{Initial} - BRET^{Final}) \times [Mpro\ sensor]} \Big/ {time}$$

where Rate = reaction velocity at a $M^{pro}$ sensor concentration

$BRET^{initial}$ and $BRET^{final}$ = initial and final BRET ratio, respectively,

[$M^{pro}$ sensor] = concentration of $M^{pro}$ sensor

time = incubation time

### Supporting Figures

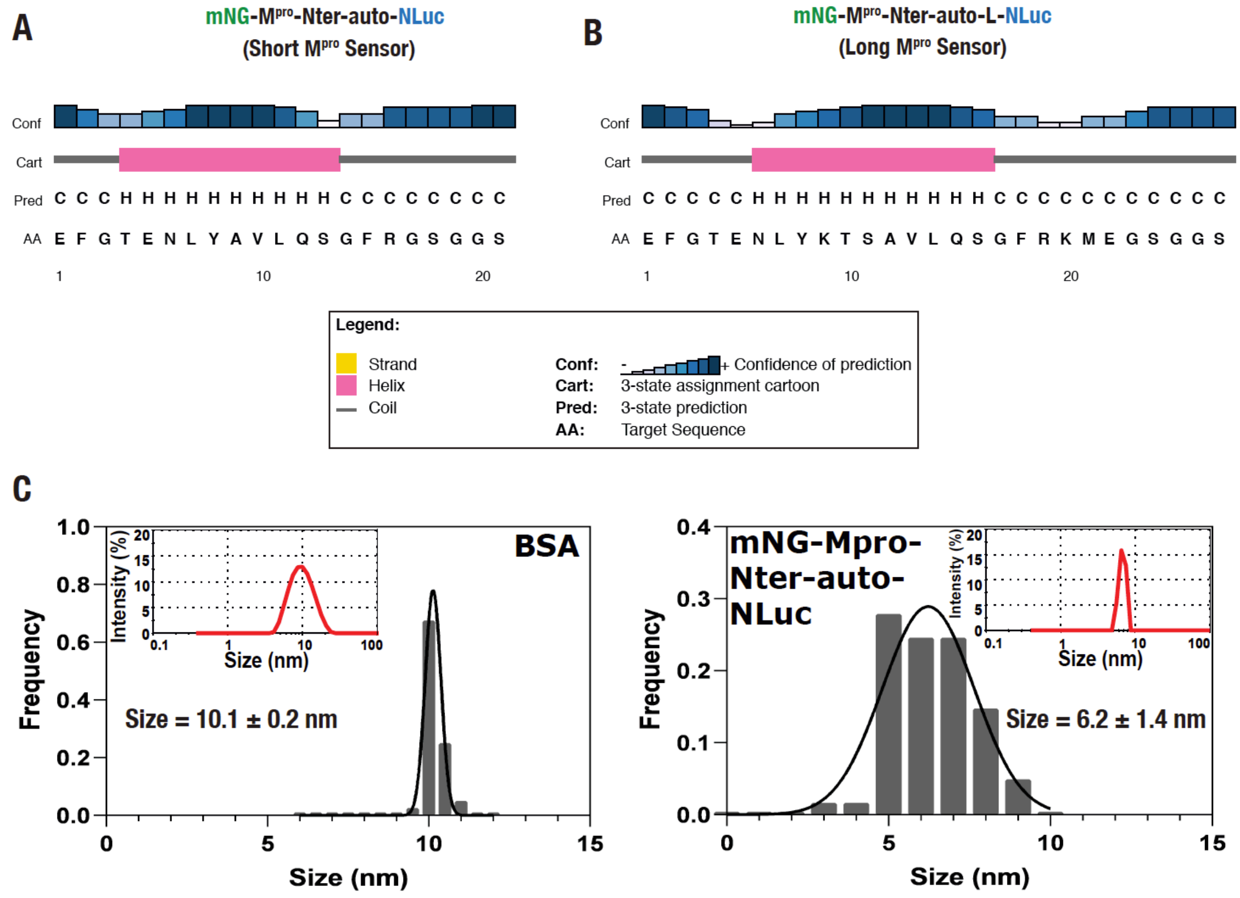


**Supp. Fig. S1.** (A,B) Secondary structure prediction of the short (A) and long (B) M$^{pro}$ BRET biosensor linkers containing M$^{pro}$ cleavage sites. (C) Graph showing frequency distribution of size (diameter, nm) of BSA (left panel) and the short M$^{pro}$ biosensor (right panel) determined from multiple ($N$ = 40 for BSA and 61 from the short M$^{pro}$ biosensor) DLS measurements. Insets in the respective graphs show a representative measurement.

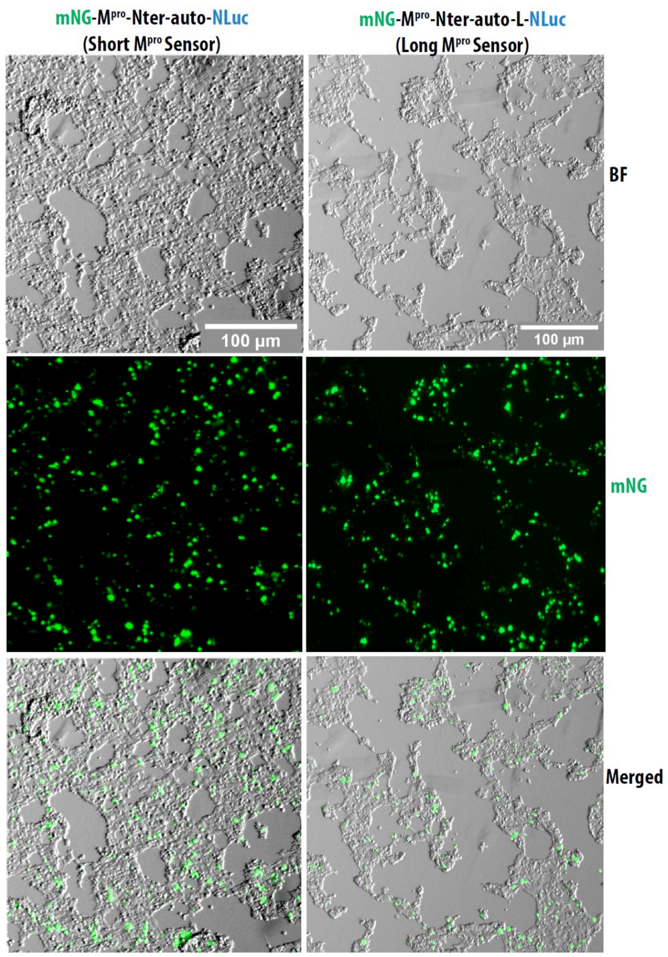


**Supp. Fig. S2. Fluorescence image of live cells showing expression of the M$^{pro}$ sensor.**

Epifluorescence images acquired using a 4× objective of HEK 293T cells transfected with either pmNG-M$^{pro}$-Nter-auto-NLuc (short; left panel) or pmNG-M$^{pro}$-Nter-auto-L-NLuc (long; right panel) plasmids showing robust expression of the sensor constructs in these cells.

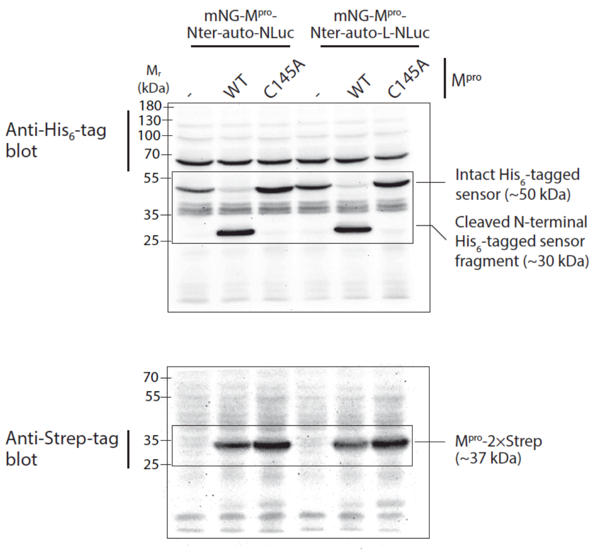


**Supp. Fig. S3. Western blot images of the M$^{pro}$ sensor constructs and M$^{pro}$.** Full western blot images used for preparing Fig. 3H top and bottom panels, respectively. Cropped regions used for preparing Fig. 3H are indicated with the rectangles in respective blots.

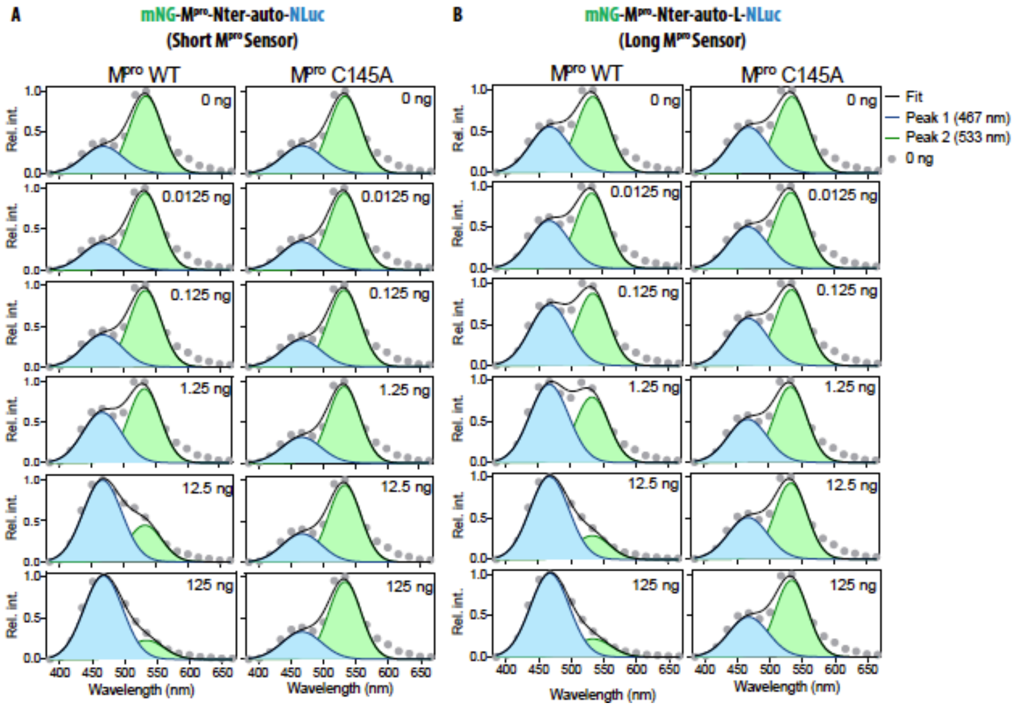

**Supp. Fig. S4. $M^{pro}$ plasmid DNA dose-dependent cleavage of the $M^{pro}$ sensors in live cells.** (A,B) Graphs showing bioluminescence spectra of the short (A) and long (B) $M^{pro}$ sensor constructs in cells expressing either the WT or C145A mutant $M^{pro}$ protease. Data were fit to a two Gaussian model reflecting mNG fluorescence and NLuc bioluminescence peaks. Note the dose-dependent cleavage of both the short (A) as well as the long (B) sensors specifically in the presence of the WT $M^{pro}$ as reflected by the reduction in the mNG peak (533 nm) of the sensor constructs. Data shown are mean ± S.D. from a representative of independent experiments performed thrice.

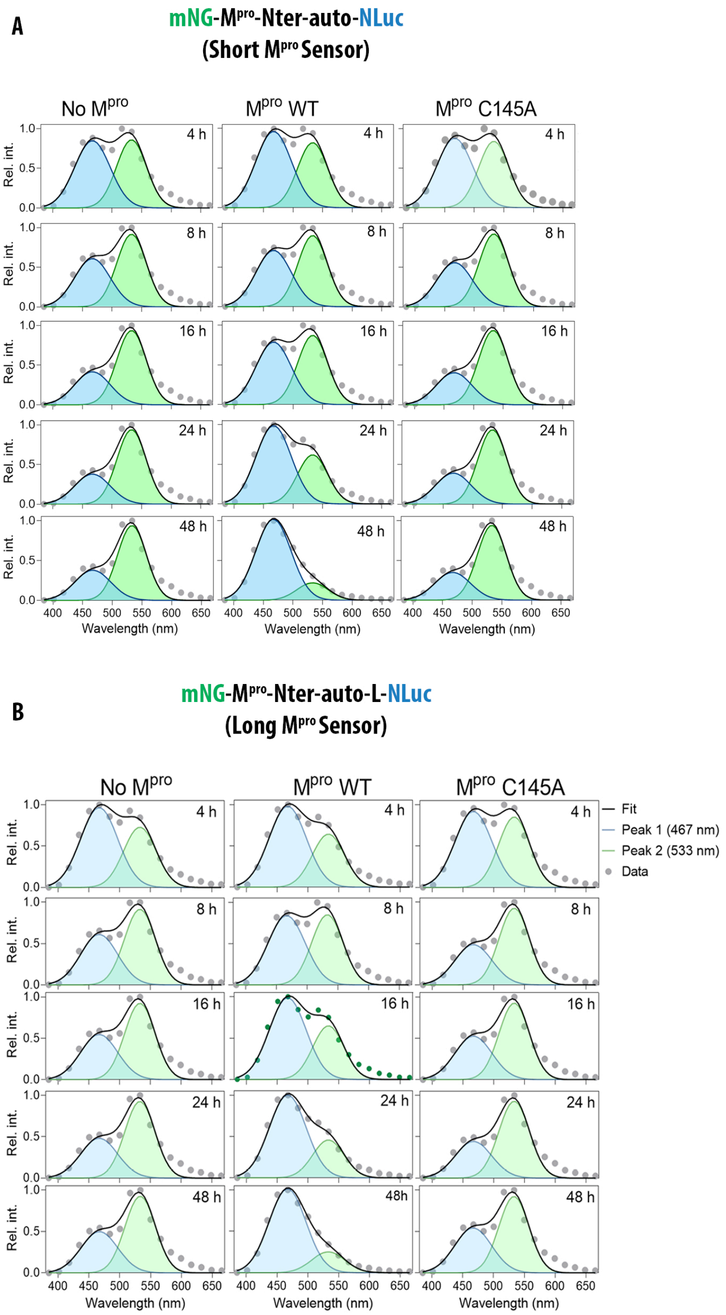


**Supp. Fig. S5. Temporal dynamics of M^pro protease activity in live cells.** (A, B) Graphs showing bioluminescence spectra of the short (A) and long (B) M^pro sensor constructs either in control cells or in cells expressing the WT or C145A mutant M^pro protease. Data were fit to a two Gaussian model reflecting mNG fluorescence and NLuc bioluminescence peaks. Note the time-dependent cleavage of short (A) and long (B) sensors specifically in the presence of the WT M^pro as reflected by the reduction in the mNG peak (533 nm) of the sensor constructs. Data shown are mean ± S.D. from a representative of independent experiments performed thrice.

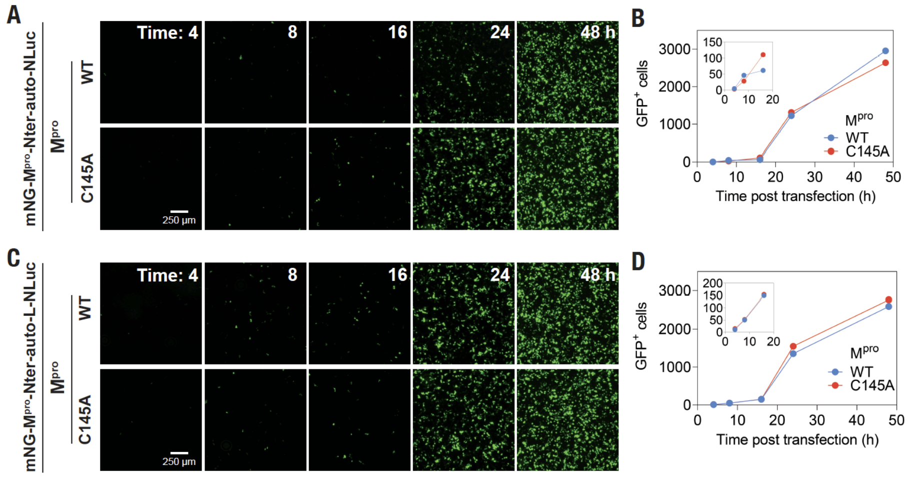


**Supp. Fig. S6. Time-dependent expression of the BRET-based M$^{pro}$ sensors.** (A,C) Epifluorescence images acquired using a 4× objective of HEK 293T cells transfected with either pmNG-M$^{pro}$-Nter-auto-NLuc (short; A) or pmNG-M$^{pro}$-Nter-auto-L-NLuc (long; C) plasmids showing a time-dependent increase in the number of cells expressing the sensors. (B,D) Graph showing time-dependent increase in GFP$^{+}$ cells after transfection with either pmNG-M$^{pro}$-Nter-auto-NLuc (short; B) or pmNG-M$^{pro}$-Nter-auto-L-NLuc (long; D) plasmids. Data shown are mean ± S.D. from a representative of independent experiments performed thrice.

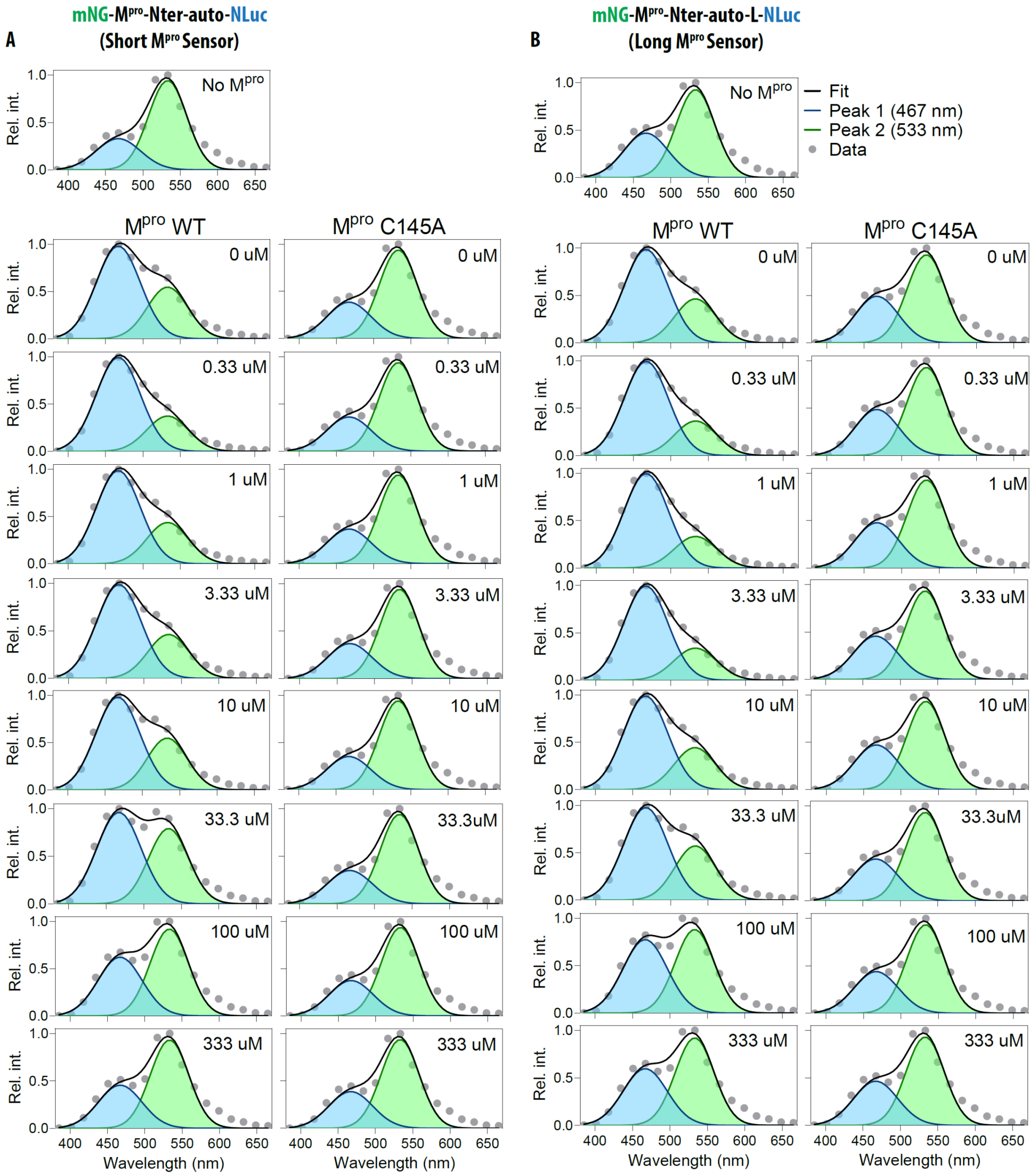


**Supp. Fig. S7. GC376-mediated M$^{pro}$ inhibition monitored in live cells.** (A,B) Graphs showing bioluminescence spectra of the short (A) and long (B) M$^{pro}$ sensor constructs in cells treated with the indicated concentrations of GC376 inhibitor in cells co-expressing either the WT or the C145A mutant M$^{pro}$ protease. The bioluminescence spectra of the No M$^{pro}$ control is also shown. Data were fit to a two Gaussian model reflecting mNG fluorescence and NLuc bioluminescence peaks. Note the dose-dependent inhibition of protease activity of WT M$^{pro}$ in cleaving both the short (A) as well as the long (B) sensors which is evident from the increased intensity of mNG peak (533 nm) of the sensor constructs. Data shown are mean ± S.D. from a representative of independent experiments performed thrice.

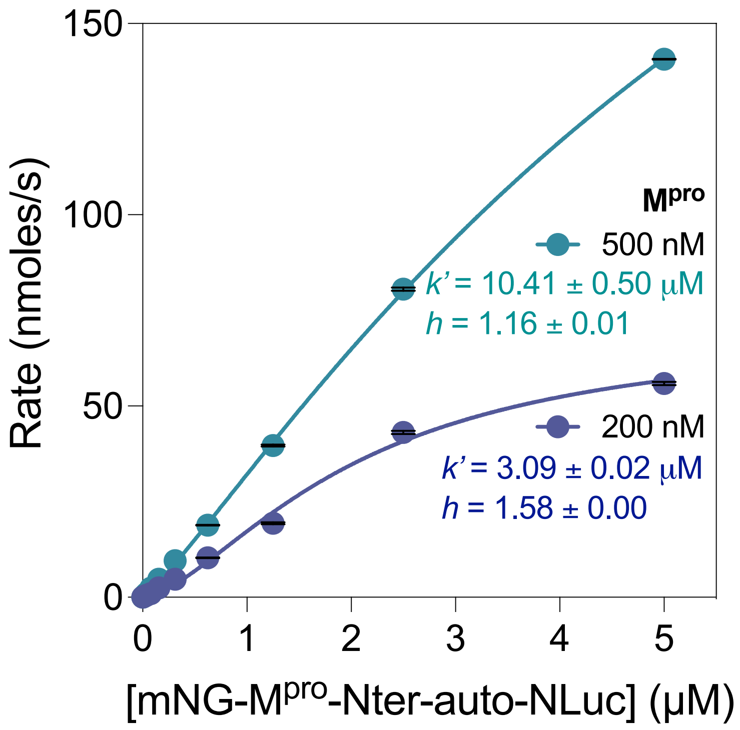


**Supp. Fig. S8. In vitro enzyme kinetics assay using the short M$^{pro}$ sensor.** Graphs showing kinetic measurements of the short M$^{pro}$ sensor cleavage in reactions containing the indicated concentrations of M$^{pro}$. Data plotted are average of four measurements ± SD and fit to the allosteric sigmoidal equation in GraphPad Prism. Note the decrease in the Hill coefficient (*h*) at 500 nM of M$^{pro}$.

### References

1. Gordon, D.E. et al. A SARS-CoV-2 protein interaction map reveals targets for drug repurposing. *Nature* (2020).
2. den Hamer, A. et al. Bright Bioluminescent BRET Sensor Proteins for Measuring Intracellular Caspase Activity. *ACS Sens* **2**, 729-734 (2017).
3. Bradford, M.M. A rapid and sensitive method for the quantitation of microgram quantities of protein utilizing the principle of protein-dye binding. *Anal Biochem* **72**, 248-54 (1976).
